# Immune activation of primary human macrophages is suppressed by the coordinated action of *Yersinia* effectors

**DOI:** 10.1101/2025.05.12.653505

**Authors:** Indra Bekere, Sören Rob, Jonas Lübbe, Susanne Kulnik, Laura Berneking, Jiabin Huang, Marie Schnapp, Björn-Philipp Diercks, Alexander Carsten, Klaus Ruckdeschel, Martin Aepfelbacher

**Affiliations:** Institute of Medical Microbiology, Virology and Hygiene, University Medical Center Hamburg-Eppendorf (UKE), Hamburg, Germany; The Calcium Signalling Group, Department of Biochemistry and Molecular Cell Biology, University Medical Center Hamburg-Eppendorf (UKE), Hamburg, Germany; Department of Bioscience, TUM School of Natural Sciences, Technical University of Munich, Lichtenbergstraße 4, 85748, Garching, Germany

## Abstract

In order to suppress the host immune response, numerous bacterial pathogens utilise a type 3 secretion system (T3SS) that injects effector proteins into host target cells. We investigated the T3SS effectors of *Yersinia enterocolitica* (Yops) for their individual and combined effects on gene expression, inflammasome formation and calcium signaling in primary human macrophages. YopP efficiently suppressed the up- and down-regulation of thousands of macrophage genes induced by the bacteria’s inflammatory stimuli. This was accompanied by parallel changes in histone 3-serine 10 phosphorylation, suggesting a higher-level regulatory mechanism. Surprisingly, YopM and YopQ counteracted selected YopP effects on gene expression, e.g., of cytokine pathways. A combination of YopP and YopQ, but not other combinations of Yops or any single Yop, reduced inflammasome formation. YopH alone blocked calcium transients in infected macrophages. We propose that the T3SS effectors of *Yersinia* antagonistically, synergistically or individually subdue major immune pathways of human macrophages to jointly suppress macrophage activation.

## Introduction

Bacterial pathogens have evolved numerous strategies that efficiently suppress the antibacterial activities of the host’s immune system in order to support their pathogenicity. Gram-negative bacteria often inactivate immune cells using specialized machineries called secretion systems, of which Type III, IV, and VI secretion systems (T3SS/T4SS/T6SS) are the most widely distributed (Costa et al., 2015; Galán & Wolf-Watz, 2006; Monjarás Feria & Valvano, 2020). These secretion systems inject effector proteins, which alone or in combination manipulate immune cell functions including cytoskeletal dynamics, e.g., to prevent phagocytosis or cell-intrinsic surveillance mechanisms, e.g., to suppress expression of inflammatory genes or block induction of cell death (Gan et al., 2021; Pinaud et al., 2018; Ruano-Gallego et al., 2021).

Pathogenic species of the Gram-negative bacteria *Yersinia*, *Escherichia coli*, *Salmonella* and *Shigella* mainly use T3SSs to inject effector proteins into human target cells (Coburn et al., 2007; Galán & Wolf-Watz, 2006). These effectors manipulate central cellular pathways through different molecular mechanisms (Aktories, 2011; Galán & Wolf-Watz, 2006). Several studies have shown that the effectors act individually but also cooperatively on host processes (Aepfelbacher et al., 2007; Malik & Bliska, 2025; Wanford et al., 2022).

Enteropathogenic *Y. enterocolitica* and *Y. pseudotuberculosis* as well as the plague agent *Y. pestis* suppress immune cell functions through seven T3SS effectors named YopE, YopT, YopH, YopO/YpkA, YopP/YopJ, YopM and YopQ/YopK (second names are *Y. pseudotuberculosis*/*Y. pestis* homologs) (Viboud & Bliska, 2005). YopE, YopT and YopO/YpkA inhibit phagocytosis and related cytoskeleton-dependent cell functions like chemotaxis by interfering with the activities of Rho GTP-binding proteins (Aepfelbacher & Wolters, 2017). YopH is a highly active tyrosine phosphatase that dephosphorylates signaling and cytoskeleton-associated proteins thereby inhibiting phagocytosis in macrophages and Ca^2+^ signaling in lymphocytes and neutrophils (Andersson et al., 1996; Andersson et al., 1999; Gerke et al., 2005). YopP/J, YopM and YopQ/K downregulate major inflammatory pathways in innate immune cells (Bliska et al., 2013; Ruckdeschel et al., 2008; Schubert et al., 2020).

YopP/J acetylates and inhibits components of NF-κB and MAPK pathways thus suppressing production of pro-inflammatory mediators and triggering cell death in macrophages (Mukherjee et al., 2006; Schubert et al., 2020). YopM associates with RSK serine/threonine kinases (p90 ribosomal S6 kinase 1; MAPKAP-K1) and PKN kinases (protein kinase N; protein kinase C related kinase/PRK) in the host cell nucleus and triggers activation of the JAK-STAT pathway to induce production of Interleukin-10 (IL-10) (Berneking et al., 2023; Berneking et al., 2016; McDonald et al., 2003). YopQ/K has been shown to regulate the injection of T3SS effectors from within host cells by an ill-defined mechanism (Dewoody et al., 2011).

Here we evaluated the individual and cooperative activities of the *Y. enterocolitica* effectors on major antibacterial immune pathways in primary human macrophages: i) Inflammatory gene expression; ii) Inflammasome activation; and iii) Increases in intracellular Ca^2+^ concentration. Primary human macrophages mirror the functions of the corresponding immune cells in the human body well and only show signs of cell death 20 hours after *Yersinia* infection. This excludes an influence of cytotoxic processes on the macrophage immune functions investigated here (Bekere et al., 2021; Ruckdeschel et al., 1997; Sarhan et al., 2018).

In earlier studies using gene microarrays, YopP was shown to efficiently suppress induction of hundreds of genes in mouse macrophages (Hoffmann et al., 2004; Sauvonnet et al., 2002). In a more recent study using RNA sequencing technologies, YopP was found to affect thousands of epigenetic histone modifications at inflammatory genes in the human macrophage genome (Bekere et al., 2021). In the past microarray study, YopM was reported to only minimally or not at all affect gene expression (Hoffmann et al., 2004; Sauvonnet et al., 2002). However, recent work demonstrated that YopM upregulates expression of multiple genes of the JAK-STAT pathway including IL-10 in human macrophages (Berneking et al., 2023; Berneking et al., 2016). Until now, an effect of YopQ/YopK on gene expression has not been reported.

YopP/J, YopM and YopQ/K have all been shown to suppress inflammasome activation in macrophages by different mechanisms. Canonical inflammasomes are cytoplasmic assemblies in which NOD-like receptors (NLRPs) acting as sensors of pathogen-associated molecular patterns (PAMPs) activate the enzyme caspase-1 via the adapter protein apoptosis-associated speck-like protein containing a CARD (ASC). Caspase-1 cleaves and activates preforms of the pore-forming molecule gasdermin D (GSDMD) and the cytokines IL-1β and IL-18, which leads to the cellular release of the cytokines through the membrane pores formed and an inflammatory cell death known as pyroptosis (Broz & Dixit, 2016).

YopP/YopJ inhibits expression of the caspase-1 substrates pro-IL1β and pro-IL18 through downregulation of NF-κB and MAPK signaling, although it may also mediate inflammasome-independent activation of caspase-1 (Ratner, Orning, Starheim, et al., 2016; Schoberle et al., 2016). YopM blocks the pyrin inflammasome, that is first activated by suppression of Rho-GTP-binding protein activity, e.g., caused by YopE, YopT or YopO/YpkA (Chung et al., 2016; Hentschke et al., 2010; Ratner, Orning, Proulx, et al., 2016). This provides a vivid example of effector triggered immunity that is counteracted by another effector (Malik & Bliska, 2025). YopQ inhibits NLRP3 and NLRC4 inflammasomes, whose activation is triggered by multiple signaling molecules or constituents of the *Yersinia* T3SS, respectively (Brodsky et al., 2010; Ratner, Orning, Proulx, et al., 2016; Zwack et al., 2015). In LPS-primed mouse macrophages, YopM and YopP/YopJ were found to cooperatively inhibit caspase-1 activation and IL1β secretion (Ratner, Orning, Starheim, et al., 2016; Schoberle et al., 2016). All studies on the *Yersinia* effects on inflammasome activity were done in mouse macrophages or mouse infection models (Bliska et al., 2013; Schubert et al., 2020). However, it is known that triggers, regulatory mechanisms and accessory proteins of inflammasomes differ between mouse- and human macrophages. E.g., activation of NLRP3 inflammasomes requires pretreatment with LPS in mouse but not in human macrophages and has a different structural basis in these species (Broz & Dixit, 2016; Xiao et al., 2023). Intracellular Ca^2+^ transients have been implicated in a plethora of macrophage functions including the control of gene transcription and inflammasome activation (Desai & Leitinger, 2014). Even though macrophages belong to the most important target cells of pathogenic yersiniae, an effect of Yops on Ca^2+^ signaling in these cells has not been reported. We report here that in primary human macrophages *Yersinia* effectors act antagonistically in regulating inflammatory gene expression (YopP vs YopM/YopQ), cooperatively in inhibiting inflammasomes (YopP and YopQ), and individually in suppressing intracellular Ca^2+^ transients (YopH).

## Results

### Global and time dependent transcriptional changes in primary human macrophages infected with *Y. enterocolitica*

First, we investigated global changes of RNA expression in primary human macrophages infected with a virulent (WA314) and a virulence plasmid cured/avirulent (WAC) *Y. enterocolitica* strain for 1.5 h and 6 h (Fig 1A). RNA expression analysis (RNA-seq) was performed using next generation sequencing (Methods) (Bekere et al., 2021; Berneking et al., 2016). Because the avirulent strain WAC lacks the T3SS, it was used to determine the effects of the PAMPs and other bacterial inflammatory stimulators. Usage of the wild-type strain WA314 allowed to determine the effect of the T3SS effectors on the inflammatory stimulator induced gene expression changes.

**Figure 1:**
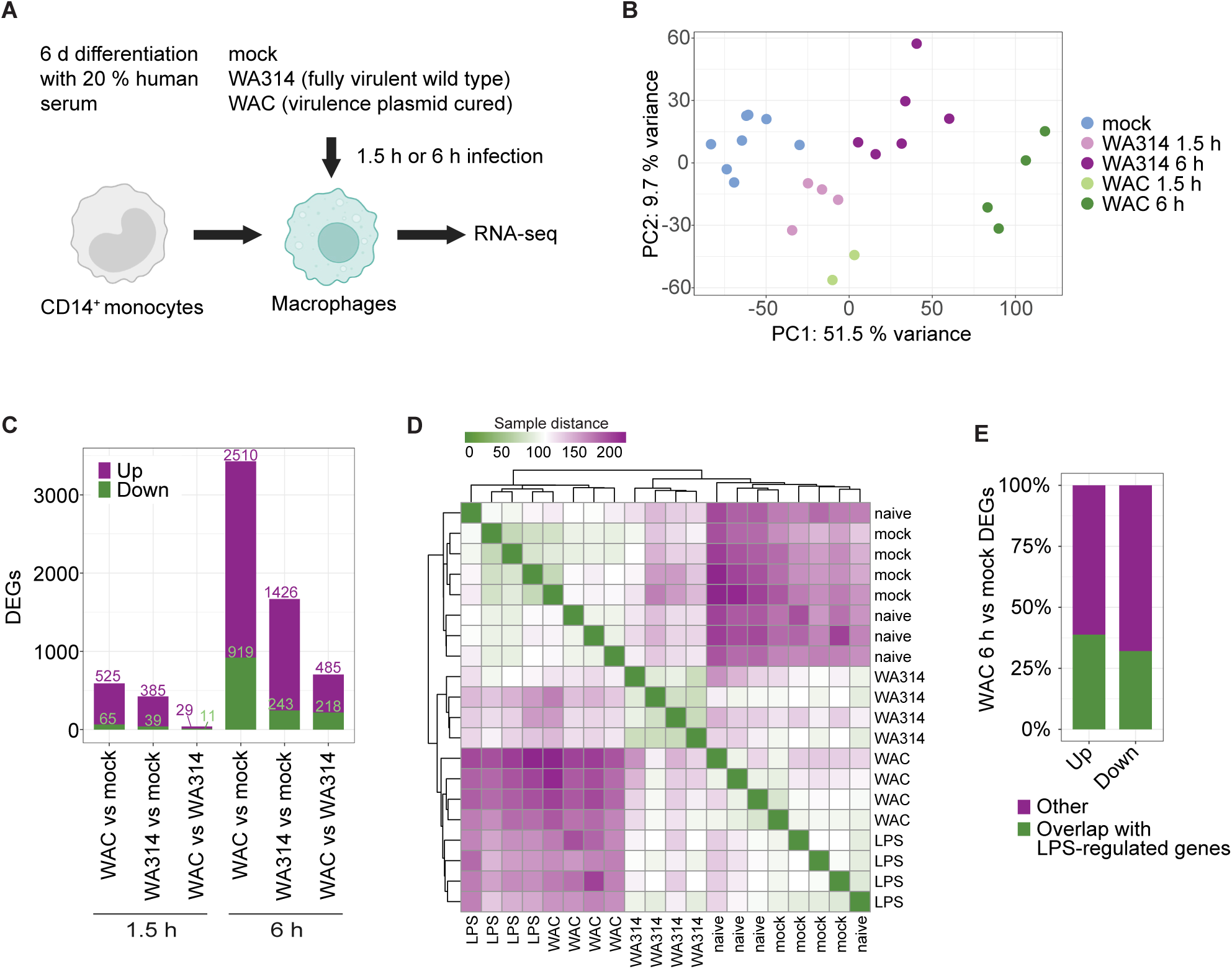
Global RNA expression analysis of *Y. enterocolitica* infected primary human macrophages. A: Experimental design. Human CD14^+^ monocytes were differentiated into macrophages by cultivation with 20 % human serum for 6 ± 1 days. Macrophages from independent donors were mock infected or infected with avirulent *Y. enterocolitica* strain WAC or wild type strain WA314 with a multiplicity-of-infection (MOI) of 100 for 1.5 h or 6 h and subjected to RNA-seq analysis. Data of all samples analyzed in this study are in Table S2. B: Principal component (PC) analysis of RNA-seq vsd gene counts of all differentially expressed genes (DEGs) identified in this study (Tables S3, S4). C: Number of up- and downregulated DEGs for indicated comparisons and infection times (Table S3). D: Heatmap representation of sample distance in human macrophages mock-infected or infected with indicated strains (6 h) combined with respective publicly available data from naive and LPS-stimulated macrophages (Novakovic et al., 2016; Park et al., 2017). E: Percentage of DEGs from WAC vs mock infected human macrophages that overlap with LPS regulated genes obtained from published data (Novakovic et al., 2016; Park et al., 2017).

Principal component analysis (PCA) of the RNA counts showed clustering of the biological replicates, which indicates high reproducibility of the experiments (Fig 1B). After 1.5 h of infection, the clusters of mock-, WAC- and WA314 infection were located closer to each other than after 6 h of infection, indicating a stronger and more pronounced transcriptional response at 6 h. Clearly separated clusters for WAC and WA314 after 6 h indicate a significant modulation of the bacterial mediator induced gene expression by the T3SS effectors (Fig 1B).

After 1.5 h of infection only slightly higher numbers of differentially expressed genes (DEGs, log2 fold change ≥ [2] and p-adjusted ≤ 0.01) were found in the WAC vs mock group as compared to the WA314 vs mock group (Fig 1C; Table S3). At this infection time, only few genes were up- and downregulated in the WAC vs WA314 group (Fig 1C). This suggests that although already after 1.5 h of infection a significant transcriptional response is caused by the *Yersinia* inflammatory mediators, only a limited T3SS effector activity is detectable at this time point.

After 6 h of infection, almost 6 times more DEGs than after 1.5 h were altogether identified (Fig 1C; Table S3). WAC vs mock showed a considerably larger number of DEGs than WA314 vs mock. This suggests that the inflammatory response of the macrophages to *Yersinia* infection is strongly repressed by the T3SS effectors after 6 h (Fig 1C). As seen in the WAC vs WA314 group, the *Yersinia* T3SS effectors altogether modulated expression of 703 genes (up- and downregulation of 485 and 218 genes, respectively; Fig 1C).

We evaluated to what extent gene expression profiles from human macrophages not infected (mock) or infected with WAC or WA314 for 6 h overlapped with naïve and lipopolysaccharide (LPS) treated macrophages (Novakovic et al., 2016; Park et al., 2017). For this we employed publicly available RNA-seq datasets from naïve and *E. coli* LPS stimulated primary human macrophages (Novakovic et al., 2016; Park et al., 2017) and compared them with our data set in a sample-to-sample distance heatmap (Fig 1D). Three groups emerged: i) naive macrophages and mock; ii) WA314 iii) WAC and LPS (Fig 1D). That the WA314 group clearly separated from the WAC/LPS group reflects the extensive modulation of the bacterial component/LPS-induced effects by the T3SS effectors (Schubert et al., 2020). On a quantitative level, 38.8 % and 32.1 % of the genes up- and down-regulated by WAC in macrophages of our study were also up- and down-regulated, respectively, by LPS in human macrophages (Fig 1E). This data indicates that the *Yersinia* induced transcriptional changes are to a large degree caused by LPS (Reinés et al., 2012). The remaining changes must then be caused by other bacterial stimulators and components, such as PAMPs, surface adhesins, constituents of the T3SS or by T3SS effector activities (Brodsky et al., 2010; Reinés et al., 2012; Zhou et al., 2018).

### Transcriptional profiles and enriched pathways in human macrophages after 1.5 h and 6 h of *Yersinia* infection

After 1.5 h of infection, 696 unique DEGs were found between mock-, WAC- and WA314 infected macrophages that could be assigned to the four clusters Early-1 to Early-4 (E1-E4; Fig 2A, B; Fig S1A; Table S4, Table S5). E1 genes were downregulated by both WAC and WA314 as compared to mock and were enriched in regulation of transcription with 17 % of the transcripts belonging to the Zinc finger C2H2-like group (Fig 2B, C, D; Fig S1B) (Lupo et al., 2013). E2 transcripts were upregulated by WA314 and WAC when compared to mock and were also enriched in transcriptional activity and included numerous central transcription factors (Fig 2B, C; Fig S1C). The transcripts in clusters E3 and E4 were strongly upregulated by WAC and this was inhibited by WA314, more so in E3 than in E4 (Fig 2A, B). E3 transcripts contain genes involved in regulation of STAT protein phosphorylation (Fig 2C, E; Fig S1D). Transcripts in the large cluster E4 were associated with inflammatory response pathways such as cytokine-cytokine receptor interaction and apoptotic process (Fig 2C; Figs S1E, F) and were consequently enriched in motifs for transcription factors RHD/NF-κB, TBP and IRF (Fig S1G). NF-κB is the main mediator of inflammatory gene expression, whereas IRFs regulate expression of secondary response genes. IRFs themselves are induced in an autocrine/paracrine fashion by type I IFNs (Kang et al., 2019; Schneider et al., 2014). We conclude that an early effect of the inflammatory mediators of *Y. enterocolitica* is to down- or upregulate transcriptional regulators. Further, E3 and E4 transcripts are primary and secondary inflammatory response genes that are induced already after 1.5 h of infection by the bacterial mediators and are already significantly (E3) or only slightly (E4) downregulated by the T3SS effectors.

**Figure 2:**
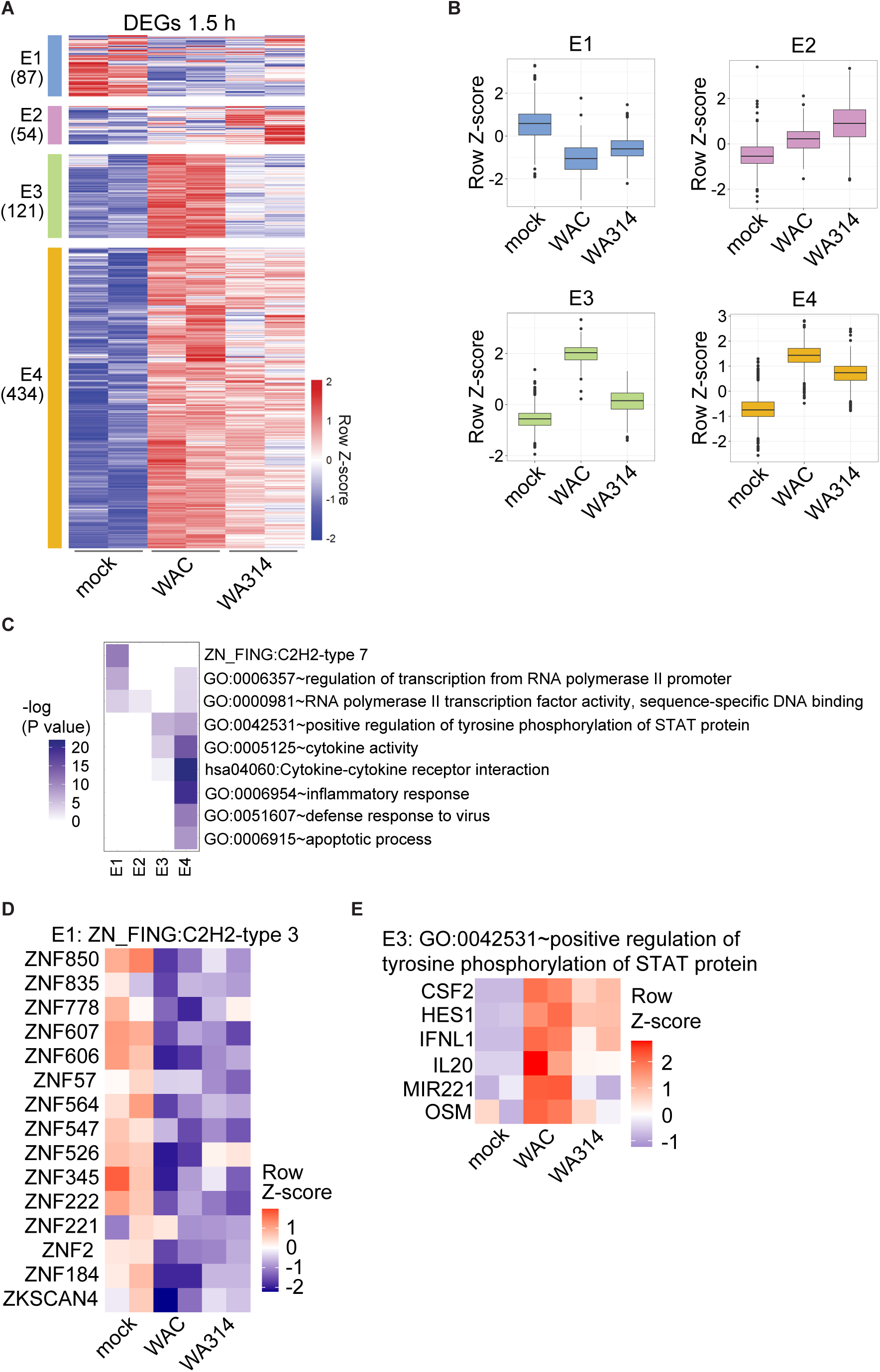
RNA expression profiles and enriched pathways in human macrophages infected for 1.5 h with *Y. enterocolitica*. A: Clustered Heatmap of DEGs (defined as log2 fold change equal or larger than 2 and adjusted P-value equal or smaller than 0.01) in comparisons between mock-, WAC- and WA314 infected macrophages after 1.5 h of infection. Gene vsd counts were row-scaled (row Z-score). 4 major clusters, E1-E4, were identified (number of genes in brackets). Two representative biological replicates are shown for each condition with all replicates shown in Fig. S1A. B: Boxplots of row scaled vsd counts for genes from clusters E1–E4 (A) showing the global expression profile in each cluster when taking into account all replicates. C: Heatmap showing –log10 transformed p-values and enriched pathways for E1-E4 (A). D: Heatmap of row scaled vsd counts for genes enriched in cluster E1 that belong to C2H2 type Zinc finger type 3. Two representative biological replicates are shown with all replicates depicted in from Fig S1B. E: Heatmap of row scaled vsd counts for genes enriched in cluster E3 that belong to the pathway positive regulation of tyrosine phosphorylation of STAT protein. Two representative biological replicates are shown with all replicates depicted in from Fig S1D.

To define the individual roles and potential interplay of the immunomodulatory T3SS effectors YopP, YopM and YopQ on regulation of gene expression in macrophages, we infected the macrophages with strains deficient in the respective Yops WA314ΔYopQ, WA314ΔYopM, WA314ΔYopP and WA314ΔYopMP for 6 h (Fig 3A). PCA of the RNA-seq data showed that the WA314ΔYopP and WA314ΔYopMP clusters located clearly separate whereas the WA314ΔYopQ and WA314ΔYopM clusters located close to the WA314 cluster (Table S3; Fig 3B). This indicates a stronger effect of YopP than of YopQ and YopM on gene transcription. Consistent with this, roughly twice as many DEGs were detected in the WA314ΔYopP vs WA314 than in the WA314ΔYopM vs WA314 comparison and only a small number of DEGs was seen in the WA314ΔYopQ vs WA314 comparison (Fig 3C). Furthermore, the number of DEGs for WA314ΔYopP vs WA314 and WA314ΔYopMP vs WA314 were about the same (Fig 3C) and the number of DEGs for WA314ΔYopP vs WA314ΔYopMP was negligible (Fig 3D). We conclude that YopP has the greatest, YopM a medium and YopQ a minor effect on inflammatory mediator induced gene transcription. Moreover, a comparison of WA314ΔYopM vs WA314ΔYopMP indicates that there is no additional effect when YopM is deleted in in the absence of YopP (Fig 3D). Therefore, the effects of the WA314ΔYopMP strain were not further included in the main results (see supplemental figures for these results).

**Figure 3:**
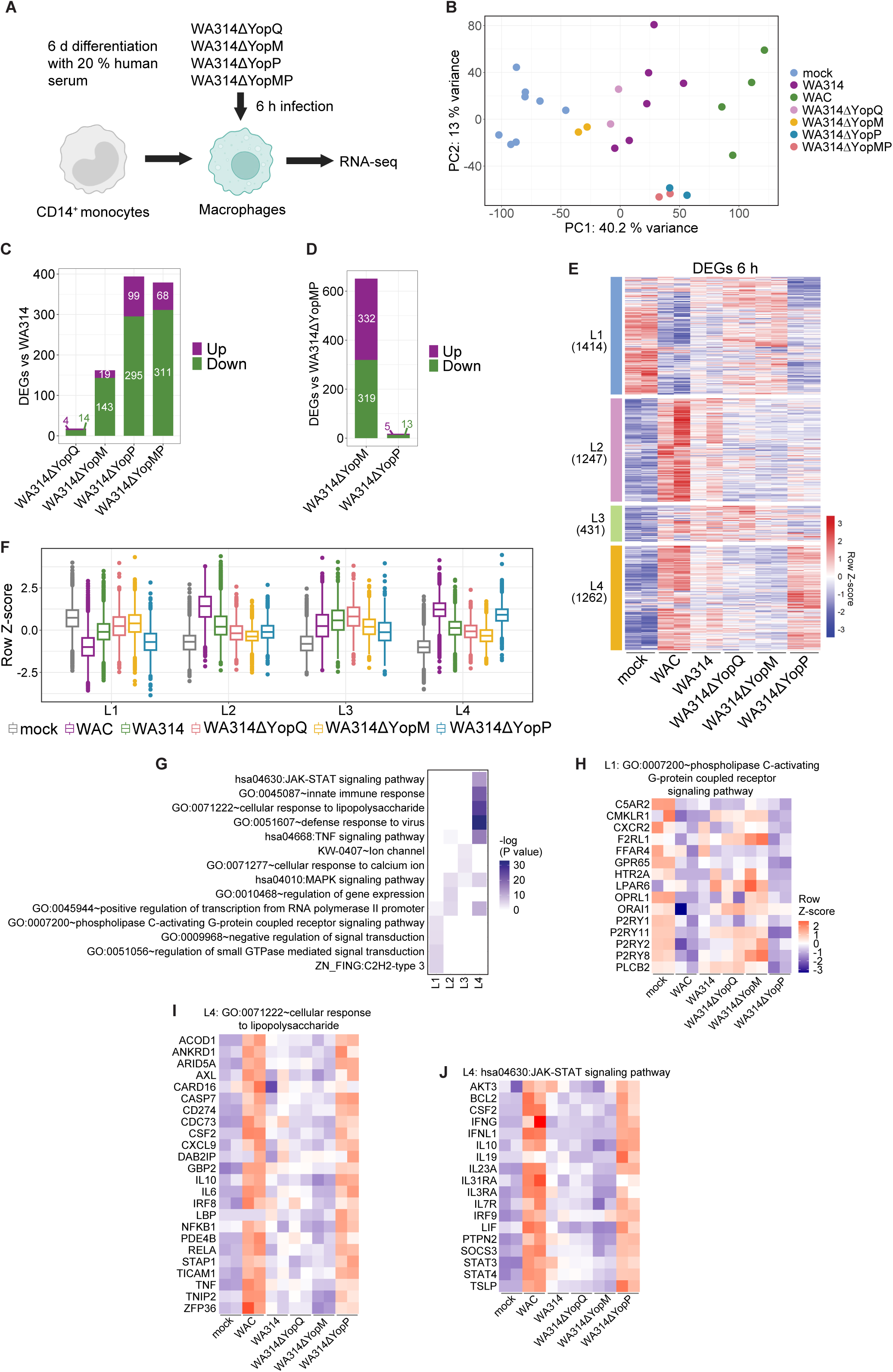
RNA expression profiles and enriched pathways in human macrophages infected for 6 h with *Y. enterocolitica*. A: Experimental setup. CD14+ monocytes were differentiated into macrophages by cultivation with 20 % human serum for 6±1 days. Macrophages from ≥ two independent donors were infected with the Yop-mutant strains WA314ΔYopM, WA314ΔYopP, WA314ΔYopMP or WA314ΔYopQ with a MOI of 100 for 6 h and subjected to RNA-seq analysis. B: Principal component (PC) analysis of vsd gene counts of all DEGs identified in this study (Tables S3, S4). C: Number of up- and downregulated DEGs in comparisons of WA314ΔYopQ, WA314ΔYopM, WA314ΔYopP and WA314ΔYopMP vs WA314 (Table S3). D: Number of up- and downregulated DEGs in comparisons between WA314ΔYopM and WA314ΔYopP vs WA314ΔYopMP (Table S3). E: Clustered heatmap of DEGs for comparisons between mock-, WAC-, WA314-, WA314ΔYopQ-, WA314ΔYopM- and WA314ΔYopP infected macrophages after 6 h. Gene vsd counts were row-scaled (row Z-score). Clusters L1–L4 were identified (number of genes in brackets). Two representative replicates are shown with all replicates and WA314ΔYopMP shown in Fig S2A. F: Boxplots of row scaled vsd counts for genes from clusters L1 – L4 (E) showing the global expression profile in each cluster when taking into account all replicates. G: Heatmap showing −log10 transformed p-values and enriched pathways for L1-L4. H: Heatmap of row scaled vsd counts for genes enriched in cluster L1 that belong to phospholipase C-activating G-protein coupled receptor signalling pathway. Two representative biological replicates are shown with all replicates depicted in from Fig S2B. I: Heatmap of row scaled vsd counts for selected genes enriched in cluster L4 that belong to cellular response to lipopolysaccharide pathway. Two representative biological replicates are shown with all replicates and genes depicted in from Fig S2I. J: Heatmap of row scaled vsd counts for selected genes enriched in cluster L4 that belong to JAK-STAT signalling pathway. Two representative biological replicates are shown with all replicates and genes depicted in from Fig S2J.

The altogether 4354 unique DEGs in macrophages infected with the different *Yersinia* strains could be assigned to 4 clusters termed Late-1 to Late-4 (L1-L4) (Fig 3E, Fig S2A, Table S6). L1 transcripts were downregulated by the bacterial mediators in WAC and this was partly prevented by WA314 in a YopP dependent manner (Fig 3F). The L1 transcripts belong to signal transduction pathways including G-protein coupled receptors, small GTPases and, as in E1 (Fig 2C), Zinc finger transcription factors (Figs 3G, H; Figs S2B, C, D). L2 transcripts were upregulated by the bacterial mediators, and this was suppressed by T3SS effectors, but YopP, YopM or YopQ did not play a significant role here (Fig 3F). The L2 transcripts are enriched in pathways of gene expression, transcription and MAPK signaling (Fig 3G, Fig S2E, F). L3 transcripts were upregulated both by bacterial mediators and the T3SS effectors YopM and YopP and contained Ca^2+^ signaling pathways (Figs 3F, G; Fig S2G). L4 transcripts, like the E3 and E4 transcripts, were upregulated by the bacterial mediators and this was suppressed by WA314 in a YopP dependent manner (Fig 3F). The L4 transcripts, like the E3 and E4 transcripts (Fig 2C), were enriched in central immune signaling pathways including cellular response to lipopolysaccharide and JAK STAT signaling that are regulated by NF-κΒ and interferon pathways (Fig 3G, I, J; Fig S2H, I, J, K).

Taken together, our data indicate that in human macrophages inflammatory *Yersinia* components up- or downregulate the expression of thousands of genes, associated with prominent pathways such as NF-κB- and interferon signaling or the regulation of gene transcription. Other pathways modulated in this context include small GTPases, JAK-STAT signaling and Ca^2+^ signaling. This extensive inflammatory response is counteracted by the combined activity of the T3SS effectors, of which YopP plays the major but not the sole role.

### YopM and YopQ antagonize YopP in the regulation of gene expression

When we analyzed the L1, L2 and L4 clusters in more detail, we noticed that the strain missing YopM and in parts also the strain missing YopQ tended to have opposite effects on gene transcription than the strain missing YopP (Fig. 3F). This phenomenon can also clearly be seen in the genes of the pathways i) phospholipase C activating G-protein coupled receptors, ii) cellular response to lipopolysaccharide and iii) JAK STAT signaling pathways (Fig 3H, I, J). To further document the antagonistic activities of YopM and YopQ vs YopP in transcriptional regulation, we performed spearman correlation analysis of the gene expression changes. The results showed a positive correlation between WAC and WA314ΔYopP and between WA314ΔYopQ and WA314ΔYopM (Fig 4A). At the same time, WAC and WA314ΔYopP showed a negative correlation versus WA314ΔYopQ and WA314ΔYopM (Fig 4A). Analysis of counterregulated genes from the RNAseq data revealed 243 DEGs in three clusters, Counterregulated-1 to Counterregulated-3 (C1-C3), that were affected in opposite directions by YopP vs YopQ and YopM (Fig 4B, C, Fig S3A, Table S7). C1 transcripts are inflammatory response genes including cytokine-cytokine receptor interaction that are downregulated by YopP. YopM and partly YopQ increased expression of these genes, thereby counteracting the YopP activity (Fig 4B, C, D; Fig S3B). Cluster C2 contains 13 genes that are upregulated by YopP but downregulated by YopQ (Fig 4B, C). These genes include calcium voltage-gated channel subunit alpha1 G (CACNA1G), Frizzled Class Receptor 4 (FZD4) involved in beta-catenin signaling and RAS protein activator like 1 (RASAL1), the latter has been described in regulating Ras-cyclic AMP pathway (Table S7) (Allen et al., 1998) (Fig 4B, C, E; Fig S3C). C3 transcripts contain genes that are downregulated by the bacterial mediators and upregulated by YopP, and this again is counteracted by YopQ and YopM (Fig 4B, C, F; Fig S3D). The C3 transcripts do not belong to a single pathway but include diverse immune signaling receptors, kinases and transcription factors (Fig 4F; Fig S3D)

**Figure 4:**
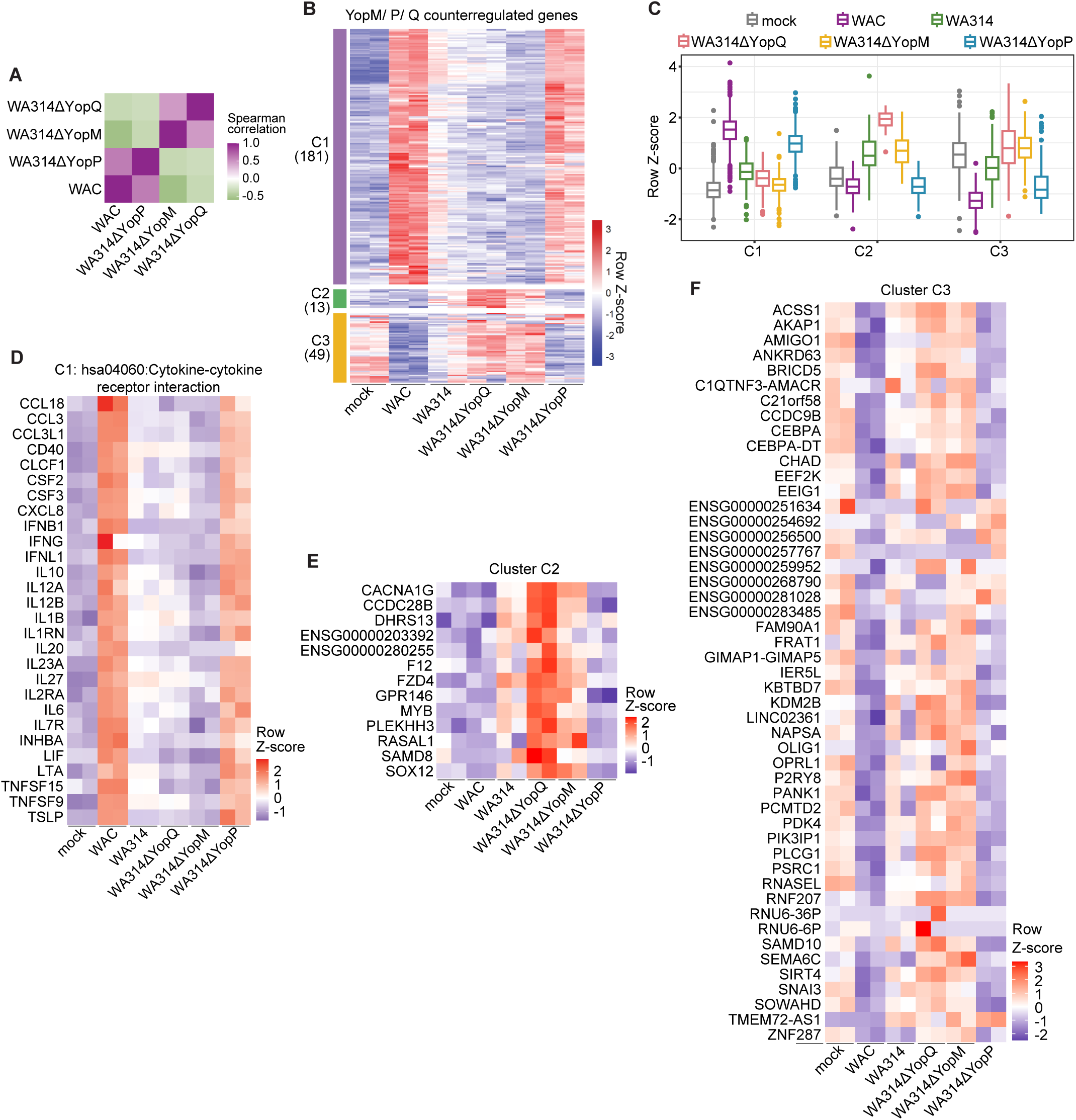
RNA expression profiles and pathways of genes counterregulated by YopP and YopM/YopQ. A: Spearman correlation heatmap representation of gene expression changes between WAC-, WA314ΔYopP-, WA314ΔYopM-, and WA314ΔYopQ infected macrophages (6 h) for WAC vs WA314 6h DEGs. B: Clustered heatmap of genes counterregulated by YopM, YopP and YopQ in comparisons between WAC- and WA314-infected macrophages (6 h). Gene vsd counts were row-scaled (row Z-score). Clusters C1 – C3 were identified (number of genes in brackets). Two representative replicates are shown, and all replicates are shown in Fig S3A. C: Boxplots of row scaled vsd counts for genes from clusters C1–C3 (B) showing the global expression profile in each cluster when taking into account all replicates. D: Heatmap of row scaled vsd counts for genes enriched in cluster C1 that belong to cytokine-cytokine receptor interaction pathway. Two representative biological replicates are shown with all replicates depicted in from Fig S3B. E: Heatmap of row scaled vsd counts for genes in cluster C2. Two representative biological replicates are shown with all replicates depicted in from Fig S3C. F: Heatmap of row scaled vsd counts for genes in cluster C3. Two representative biological replicates are shown with all replicates depicted in from Fig S3D.

We conclude that part of the YopP effects on gene transcription are systematically counteracted by YopQ and YopM. This counteraction involves key immune signaling pathways of macrophages and therefore are likely to be of cell biological relevance.

### *Yersinia* effectors suppress phosphorylation of H3S10 at heterochromatin in macrophages

The profound effects of *Yersinia* on macrophage gene expression prompted us to search for higher-level regulatory mechanisms potentially underlying this phenomenon. Bacterial inflammatory mediators activate NF-κB and MAP-kinase (MAPK) signaling pathways which regulate gene expression through a plethora of mechanisms (Liu et al., 2017; Whitmarsh, 2007). Notably, MAPK signaling has been shown to induce phosphorylation of histone-3 at serine 10 (H3S10ph), an epigenetic modification that primes chromatin for deposition of other modifications or factors that regulate gene expression (Dong & Hamon, 2020; Prigent & Dimitrov, 2003; Sawicka & Seiser, 2012). H3S10ph can thereby control gene expression profiles involving hundreds of genes (Prigent & Dimitrov, 2003; Sawicka & Seiser, 2012). Western Blot analysis revealed that H3S10ph was virtually absent in mock infected macrophages whereas both WAC and WA314 induced H3S10ph already after 10 min of infection (Fig 5A). From 30 to 90 min, WAC infected cells showed high levels of H3S10ph, whereas in WA314-infected cells H3S10ph was reduced compared to WAC and even absent after 90 min (Fig. 5A). The suppression of H3S10ph by WA314 was confirmed by two different anti-H3S10ph antibodies (Fig S4A). This indicates that H3S10ph is induced rapidly by the bacterial mediators and is thereafter suppressed by the T3SS effectors. These kinetics also correlate well with the stimulation of inflammatory gene expression by both, WAC and WA314 after 1.5 h of infection and with suppression of gene expression by WA314 after 6 h of infection (Figs 2 and 3). Pretreatment with MAPK inhibitors (PD+SB) and an NF-κB inhibitor (TPCA) suppressed H3S10ph after WAC infection, indicating that H3S10ph is induced through MAPK and NF-κB mediated pathways (Fig 5B).

**Figure 5.**
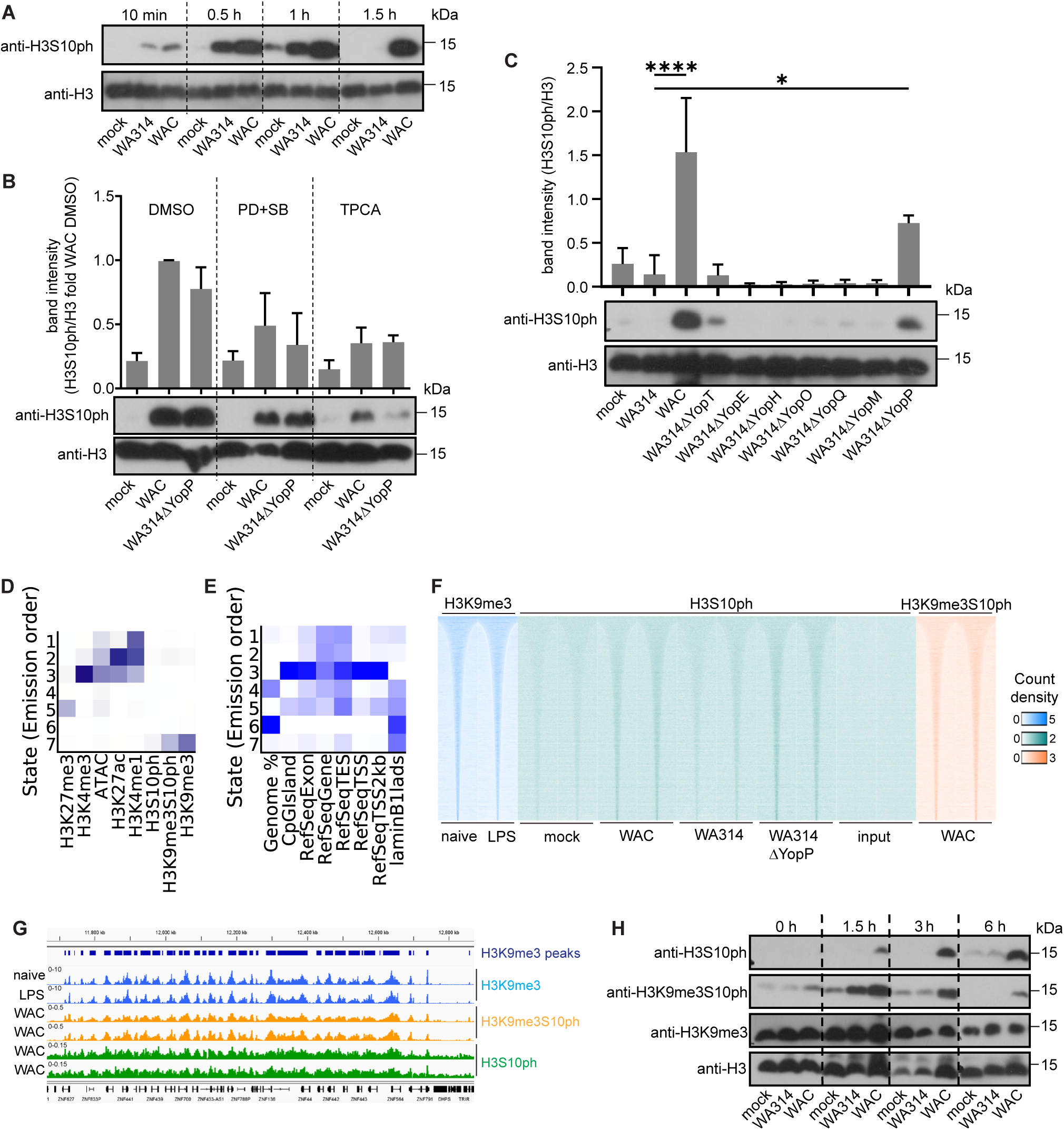
*Yersinia* effectors suppress phosphorylation of H3S10 at heterochromatin regions in human macrophages. A: Western blot of H3S10ph levels in macrophages that were mock infected or infected with WA314 or WAC for 10 min to 1.5 h with MOI of 100. Histone-3 (H3) bands serve as loading control. B: Bar graph and representative Western blot (bottom) of H3S10ph levels in macrophages pretreated with solvent (DMSO), MAPK inhibitors (PD+SB) or NF-κB inhibitor (TPCA) for 30-60 min followed by infection with indicated strains for 3 h with MOI of 100. H3 bands serve as loading control. Bars show mean and standard deviation from three independent experiments. C: Bar graph and representative Western blot (bottom) of H3S10ph levels in macrophages mock infected or infected with indicated strains for 3 h with MOI of 100. H3 bands serve as loading control. Bars show mean and standard deviation from three independent experiments. P-value calculations were performed using one-way Anova. * P ≤ 0.05, **** P ≤ 0.0001. D: Heatmap showing ChromHMM analysis assigning different chromatin modifications or open chromatin regions from ATAC-seq (columns) to different chromatin states (1-7; rows). The darker blue color corresponds to a greater probability of observing the mark in the state. Heatmap shows that H310ph, H3K9me3 and H3K9me3S10ph regions are found in state 7 described by closed chromatin (no signal from ATAC-seq). H3S10ph ChIP-seq showed very low enrichment levels. E: Heatmap from ChromHMM analysis showing enrichment for various genomic annotations in different states. The analysis shows that state 7 containing H310ph, H3K9me3 and H3K9me3S10ph regions (D) is enriched for heterochromatin characterized by lamin B1 lamin-associated domains (lads). F: Heatmap showing H3S10ph ChIP-seq signal (green) from macrophages mock infected or infected with the indicated *Yersinia* strains or from input control at H3K9me3 peaks from naïve and LPS-treated macrophages from publicly available dataset (blue) (Novakovic et al., 2016). H3K9me3S10ph ChIP-seq from WAC-infected macrophages is depicted in orange. Rows are genomic regions from −10 to +10 kb around the centre of the analyzed regions. n = 64419 G: Peak tracks of ChIP-seq tag densities at H3K9me3 peaks (dark blue bars) for H3K9me3 (blue), H3K9me3S10ph (yellow) and H3S10ph (green) for WAC-infected macrophages and naïve- and LPS-stimulated macrophages from publicly available dataset (Novakovic et al., 2016). H: Western blot showing H3S10ph-, H3K9me3S10ph- and H3K9me3 levels in macrophages mock infected or infected with WA314 or WAC for 0 to 6h with MOI of 100. H3 bands serve as loading control.

To investigate which of the effectors may suppress H3S10ph, the macrophages were infected with strains lacking each of the effectors individually (Fig 5C). *Yersinia* mutants lacking YopT, -E, -H, -O, -Q or -M suppressed H3S10ph to the same degree as WA314, indicating that none of these effectors alone is capable of inhibiting H3S10ph (Fig 5C). In comparison, WA314ΔYopP inhibited H3S10ph about half as effectively as WA314 (Fig 5C). This suggests that YopP plays the major role in suppressing H3S10ph and that the remaining inhibition of H3S10ph is due to a combined effect of the other Yops. Treatment with the MAPK- and NF-κB inhibitors when compared to DMSO control reduced H3S10ph in WA314ΔYopP infected cells, consistent with the idea that YopP inhibits H3S10ph through inhibition of MAPK and NF-κB signaling (Fig 5B).

In the next step we performed H3S10ph chromatin immunoprecipitation and sequencing (ChIP-seq) to analyze at which chromatin regions H3S10ph occurs. The macrophages were mock-infected or infected with WAC, WA314 or WA314ΔYopP for 3 h followed by H3S10ph ChIP-seq (Methods). We performed chromatin state discovery and characterization with ChromHMM analysis (Ernst & Kellis, 2012, 2017) to test whether H3S10ph is localized at active/open chromatin and may associate with inflammatory genes or inactive/closed chromatin states. For this, we integrated our own data set from H3K4me3, H3K27me3, H3K4me1 and H3K27ac ChIP-seq in *Yersinia*-infected primary human macrophages (Bekere et al., 2021) and publicly available data sets from H3K9me3 ChIP-seq and ATAC-seq in primary human macrophages (Novakovic et al., 2016) H3S10ph showed weak enrichment but notably, was detected in regions marked by repressive heterochromatin mark H3K9me3 (Figs 5D, F, G). The H3K9me3 mark is found in lamin B1 lamin-associated domains (LADs) that signify inactive and closed chromatin (Fig 5E). H3S10ph was not associated with active chromatin regions containing modifications associated with active gene transcription such as H3K4me3 and H3K27ac, enhancer regions marked with H3K4me1 or open chromatin regions from ATAC-seq data (Fig 5D, E). We next employed an antibody that exclusively recognizes the double chromatin modification H3K9me3/H3S10ph but not the individual H3K9me3 and H3S10ph modifications. Western blot analysis revealed that the combined H3K9me3/H3S10ph modification increased in parallel to the H3S10ph modification in WAC treated cells and was strongly inhibited like the H3S10ph modification in the WA314 treated cells from 3 h to 6 h of infection (Fig 5H). In contrast, the H3K9me3 mark alone did not show these kinetics (Fig 5H). ChIP-seq analysis using the H3K9me3/H3S10ph antibody in the WAC infected macrophages showed a better enrichment in the H3K9me3 marked regions than when the H3S10ph antibody was used (Fig 5D, F, G). We conclude that the massive initiation of gene transcription by the *Yersinia* mediators is associated with a strongly increased deposition of H3S10ph near the repressive chromatin mark H3K9me3, which may prepare/prime inactive chromatin regions for access by transcription-promoting factors (Wong et al., 2024). Conversely, inhibition of gene transcription by YopP and the cooperative action of the remaining Yops is associated with a significantly reduced amount of H3S10ph at the repressive chromatin mark, which most likely inhibits gene transcription on a global level.

### *Yersinia* effectors YopQ and YopP cooperate to downregulate inflammasome formation in human macrophages

It has been reported that YopQ/YopK suppresses the activation of the NLRP3 inflammasome induced by the T3SS translocation pores and that YopM inhibits the activation of the pyrin inflammasome induced by the inactivation of Rho-GTP-binding proteins (Brodsky et al., 2010; Malik & Bliska, 2020; Ratner, Orning, Proulx, et al., 2016). These results were obtained in bone marrow derived mouse macrophages and no data on the individual or combined effects of *Yersinia* T3SS effectors in primary human macrophages have been reported. To investigate formation of inflammasomes in primary human macrophages, we infected the cells with strain WAC(pT3SS), a WAC strain harboring and overproducing the T3SS including translocation pores but lacking the Yop effectors (Heesemann & Laufs, 1983). WAC(pT3SS) infection produced time and dose dependent ASC specks in the macrophages (Fig 6A, B). These specks contain NLRP3-GFP and Caspase-1 and therefore qualify as clusters of inflammasomes (Fig 6A) (Nagar et al., 2023). Macrophages not infected (mock) or infected with avirulent WAC, wild type WA314, or with wild type strains containing single deletions of YopM, YopP or YopQ showed only a minimal ASC speck formation (Fig 6C). Altogether this data shows that i) the *Yersinia* T3SS structural components are strong activators of inflammasome formation, ii) *Yersinia* inflammatory mediators alone (like LPS) do not cause formation of inflammasomes, iii) WA314 through its Yops completely blocks inflammasome formation triggered by the T3SS components, and iv) the removal of YopQ, YopM or YopP alone is not sufficient to reverse the blockage of inflammasome formation. To find our whether Yops cooperate to inhibit inflammasome formation, we infected the macrophages with the double mutants lacking YopM /YopP, YopM/YopQ and YopP/YopQ and the triple mutant lacking YopM/YopP/YopQ. The YopP/YopQ and YopM/YopP/YopQ mutants showed significant and equal increases in inflammasome formation whereas the other mutants had no effect (Fig 6C). That YopM had no effect on inflammasome formation in primary human macrophages, either alone or in combination with YopQ and/or YopP, is in contrast to mouse macrophages, in which YopM plays an important role in blocking the pyrin inflammasome (Malik & Bliska, 2020). Overall, these data suggest that YopP and YopQ must work together to inhibit inflammasome formation in primary human macrophages.

**Figure 6.**
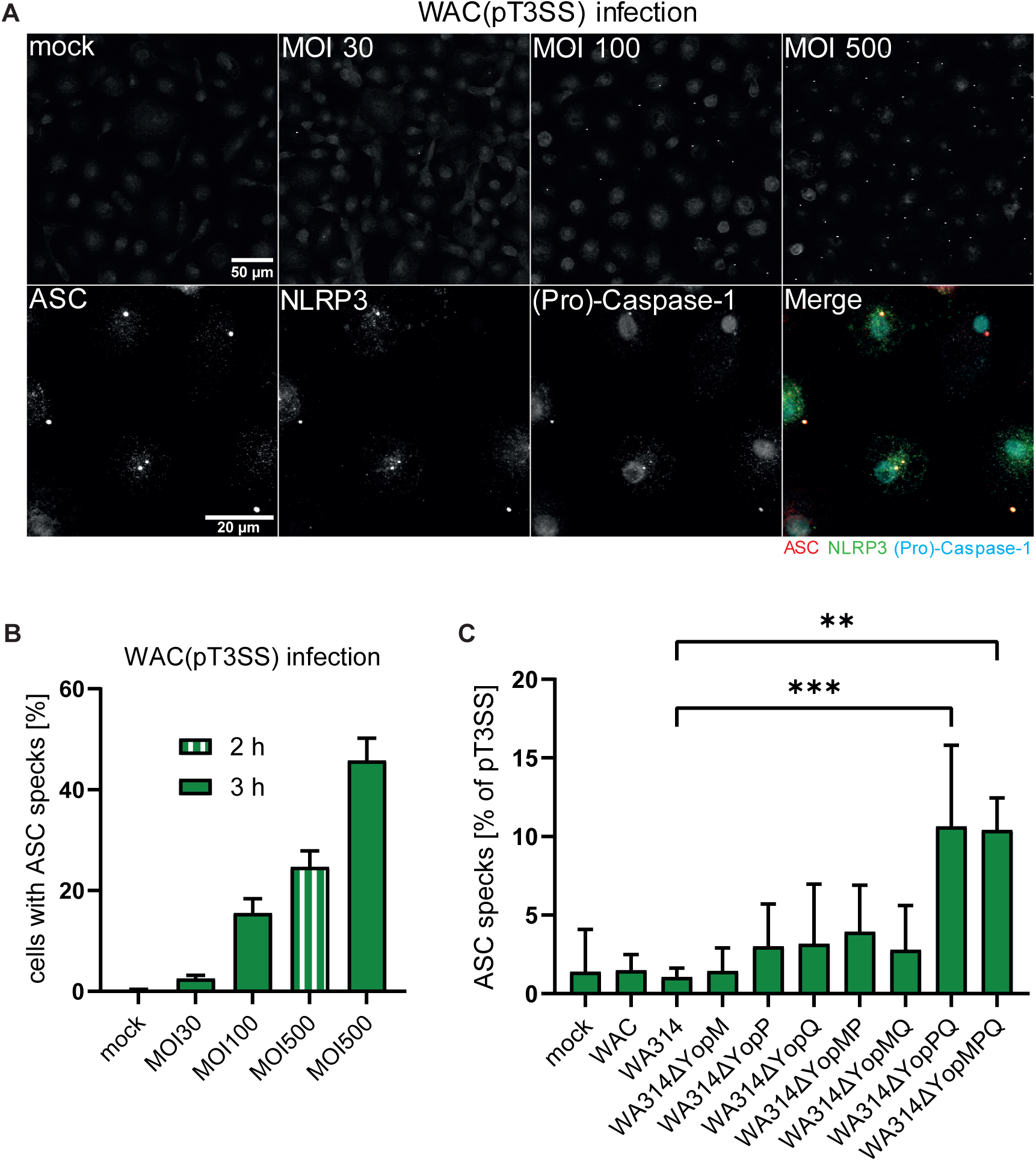
*Yersinia* effectors cooperate to downregulate inflammasome formation in human macrophages. A) (Upper row) Human macrophages form inflammasomes upon infection with a *Yersinia* strain that overproduces translocation pores. Macrophages were mock-infected or infected with indicated MOIs of strain WAC(pT3SS) for 3 h and immunostained for endogenous ASC. (Bottom row) NLRP3-eGFP transfected human macrophages were infected with WAC(pT3SS) at a MOI of 500 for 2 h and immunostained for endogenous ASC and (Pro)-Caspase-1. Merge shows overlay of ASC (red), NLRP3 (green) and (Pro)-Caspase-1 (blue). B) Dose and time dependency of ASC speck formation in human macrophages infected with indicated MOIs of WAC(pT3SS) for 3 h or 2 h. The 2 h value is shown because values in (C) were normalized to it. Bars represent mean and standard deviation from at least two biological replicates with 600 cells counted per replicate. C) Effect of individual or combined *Yersinia* effector mutants on NLRP3 inflammasome formation in human macrophages. Macrophages were mock-infected or infected with indicated strains at a MOI of 500 for 2 h. Bars represent mean and standard deviation of at least two replicates with 600 cells counted per replicate. Values were normalized to values of WAC(pT3SS) infected macrophages (MOI 500, 2 h; C) in the respective experiment. Significance was calculated using an ordinary one-way ANOVA. *: p-value ≤ 0.05.

### *Yersinia* effector YopH suppresses Ca^2+^ spikes in human macrophages

Changes in the intracellular Ca^2+^-concentration are a universal regulator of macrophage immune activities, including gene expression and cytokine production (Desai & Leitinger, 2014). Although *Y. pseudotuberculosis* has been described to affect intracellular Ca^2+^ fluctuations via the effector YopH in human neutrophils (Andersson et al., 1999; Rolan et al., 2013), there are no reports on how Ca^2+^ responses are altered by pathogenic *Y. enterocolitica* in macrophages. Ca^2+^-imaging of mock-infected macrophages revealed that within a period of 120 min Ca^2+^ fluctuations accumulated to 7.6 Ca^2+^ spikes/cell with a mean increase of 54 nM (Fig. 7A, B; Fig. S5A, B). In WAC-infected cells, number and mean increase of Ca^2+^ spikes were significantly elevated compared to mock (34 Ca^2+^ spikes/cell; increase: 157 nM) (Fig 7A, B, Fig S5B). This suggests stimulation of Ca^2+^ signaling by LPS or other activators such as Invasin. WA314-infected cells showed much lowered number and mean increase of Ca^2+^ spikes (1.4 Ca^2+^ spikes/cell; increase: 36.4 nM) suggesting that one or more effectors suppress both, base-line and *Yersinia* mediator induced Ca^2+^ fluctuations (Fig 7A, B; Fig S5A, B). Infection of the macrophages with single mutants of each of the seven Yops showed that only infection with the YopH-deficient strain resulted in Ca^2+^ spikes that were similar in number and increase to the spikes in WAC infected cells (25.9 Ca^2+^ spikes/cell; increase: 66.8 nM) (Fig. 7A, B; Fig. S5B; movie S1). We conclude that baseline and *Yersinia* stimulated Ca^2+^ signaling in human macrophages is specifically and selectively abolished by YopH.

**Figure 7.**
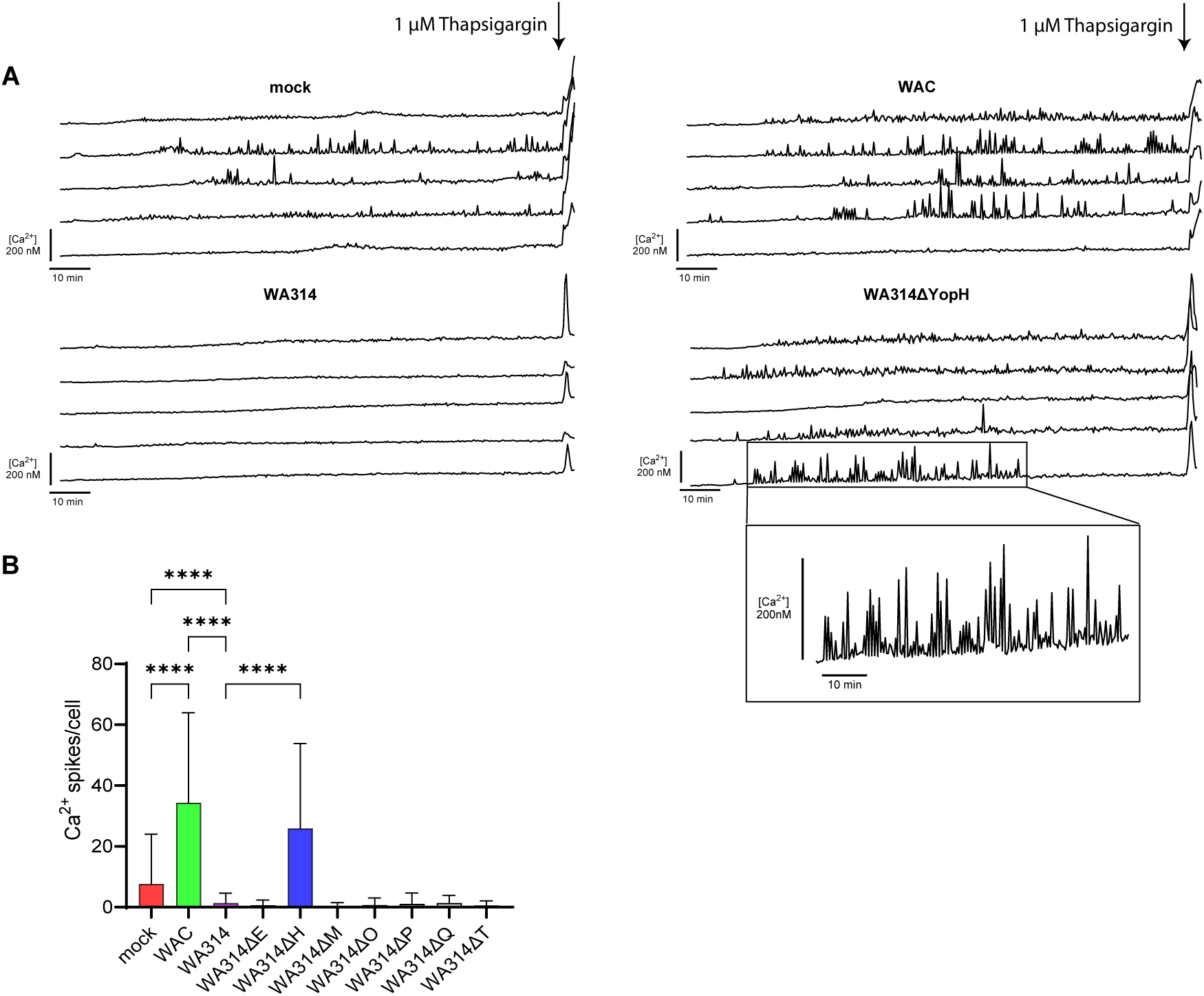
*Yersinia* effector YopH blocks rises in intracellular Ca^2+^-concentration in human macrophages. A: Five representative single-cell Ca^2+^-concentration changes in macrophages mock-infected or infected with each of the indicated *Yersinia* strains. Human macrophages were not infected (mock) or infected with indicated strains with a MOI of 50 for 2 h and loaded with Cal520-AM and FuraRed-AM. Cells were imaged with 3 frames per minute for 2 h. Stimulation with 1 µM Thapsigargin at 2 h post infection (hpi) was used as a positive control. B: Quantification of Ca^2+^ spikes/cell accumulated until 2 hpi. Ca^2+^ spikes were defined as ΔF_em1_>20% (Methods). Data represent mean and standard deviation of n=3 independent experiments with altogether 238-673 cells evaluated for each condition. For statistical analysis ordinary one-way ANOVA with multiple comparisons was used. ****P≤0.0001.

## Discussion

The strategic cooperation between bacterial virulence factors was referred to as a stratagem several decades ago (Heesemann et al., 2006; Jann & Jann, 1987). Numerous reports meanwhile indicate that the activities of the different effectors of a pathogen can be linked in a variety of ways (Sanchez-Garrido et al., 2022). A recent article investigated the network of T3SS effectors of *Citrobacter rodentium* in mouse pathogenicity (Ruano-Gallego et al., 2021). One mutant strain that lacked 19 of the 31 *C. rodentium* effectors was still virulent, indicating considerable redundancy between the effectors in this pathogen. The remaining 12 effectors formed a robust network and could not be further contracted without compromising virulence (Ruano-Gallego et al., 2021). In comparison, pathogenic *Y. enterocolitica* spp. contain only seven effectors, and infection of mice suggest that each of the five effectors - YopP, YopQ, YopM, YopE and YopH - is essential for the maintenance of full pathogenicity (Trülzsch et al., 2004). Thus, most of the *Yersinia* effectors already fulfil unique and non-redundant functions during infection. Most studies investigating Yersinia T3SS effectors have focused on the activity of individual effectors in transformed cell lines or mouse macrophages. To obtain more detailed information on how key immune functions of human macrophages are jointly altered by the Yersinia effectors, we here tested their individual and especially combined effects on the expression of inflammatory genes, the activation of inflammasomes and changes in intracellular Ca^2+^ concentration. YopP had the most profound effect on the inflammatory mediator induced changes in gene transcription. It suppressed the up- or downregulation of altogether > 2500 genes most certainly by virtue of its inhibition of NF-κB- and MAP-kinase signalling. As a potential higher-level regulatory mechanism of gene transcription in infected macrophages, we investigated the level of phosphorylated serine-10 in histone-3 (H3S10ph) which was shown to be induced by LPS in mouse macrophages (Josefowicz et al., 2016) and is efficiently subverted by pathogenic bacteria *Listeria monocytogenes*, *Streptococcus pneumoniae* and *Pseudomonas aeruginosa* (Dong et al., 2020; Dortet et al., 2018; Hamon & Cossart, 2011). The Yersinia inflammatory mediators induced a NF-κB- and MAP-kinase-dependent increase followed by a YopP-mediated decrease of H3S10ph. H3S10ph colocalized with the repressive histone mark H3K9me3 in the macrophage genome, suggesting that it regulates the accessibility of genes to transcriptional activators (Sawicka & Seiser, 2012; Yamamoto et al., 2003). Phosphorylation of H3S10 at H3K9me3 is a mechanism known during mitosis as “phosphor-methyl switch” regulating HP1 dissociation and cell division (Fischle et al., 2005; Hirota et al., 2005), while phosphorylation of H3S10 at H3K9me2 regulates gene expression upon growth factor stimulation (Wong et al., 2024). We show that in *Yersinia*-infected primary human macrophages H3K9me3S10ph is likely involved in rapid induction of transcriptional response by inflammatory mediators of bacteria and inhibition of H3S10ph by YopP contributes to its overwhelming negative effect on gene transcription.

Surprisingly, we found that YopM and partly also YopQ counteracted selected effects of YopP on gene transcription, e.g., of cytokine signaling pathways. Previous studies have already shown that YopM can reverse the YopP-mediated downregulation of IL-10 production. To do so YopM caused activation RSK kinases in the host cell nucleus and nuclear translocation of the transcription factor Stat3 (Berneking et al., 2023; Berneking et al., 2016). The additional > 200 genes identified here as oppositely regulated between YopP and YopM/YopQ include not only a large number of cytokines but also components of other inflammatory signalling pathways. How the opposing regulation of these genes eventually contributes to the pathogenicity of the bacterium remains unclear at the moment, but it can be speculated that YopM reverses exactly those activities of YopP that would otherwise hinder the infection progression. For example, reversal of the YopP-inhibited expression of the immunosuppressive cytokine IL-10 and JAK-STAT signaling genes may be beneficial for the pathogen (Berneking et al., 2023; Duell et al., 2012). Furthermore, YopM may balance expression of cell survival and inflammatory genes, that are suppressed by YopP/J thus regulating induction of caspase-8-dependent cell death that is beneficial for the host (Monack et al., 1997; Zhang et al., 2005). Another finding of our study is that although YopQ by itself had no major influence on gene expression, a group of signaling genes whose products are involved in Ca^2+^ -, beta-catenin- and RAS signaling are downregulated by YopQ. This result could be a first step towards the elucidation of the molecular mode of action of YopQ.

When we investigated the suppression of inflammasome/speck formation by the *Y. enterocolitica* effectors, we found that single knockouts of each of the three effectors - YopM, YopQ and YopP – had no effect in human macrophages. Only after infection with the combined YopP/YopQ-deficient strains suppression of speck formation was significantly reduced. We conclude that, as opposed to the situation in mouse macrophages, YopM neither alone nor in combination with YopP or YopQ affects inflammasome/speck formation in human macrophages. A possible explanation for this could be the strongly altered expression of Rho-GTPase genes, which was previously described in human macrophages infected with *Y. enterocolitica* (Bekere et al., 2021). One may speculate that alternative Rho GTP binding proteins produced in the macrophages during infection are insensitive to YopE and YopT and could take over their function in keeping the pyrin inflammasome inactive. The finding that a combination of YopP and YopQ was required to inhibit inflammasome/speck formation may be explained by the fact that in the human macrophages LPS-triggered expression of inflammasome constituents and NLRP3-mediated inflammasome activation must be inhibited in parallel by YopP and YopQ, respectively (Kelley et al., 2019). Future studies should investigate these intriguing possibilities. Finally, as was shown before in neutrophils, calcium signaling in form of repeated fluctuations of intracellular Ca^2+^ concentration in the *Yersinia* infected macrophages was blocked completely by YopH and none of the other effectors had any effect on it. In summary, we propose that Yersinia effectors act antagonistically, cooperatively and individually on key immunoregulatory pathways in human macrophages to ultimately suppress the activation state of these immune cells.

## Materials and Methods

### Bacterial strains

*Yersinia enterocolitica* strains used in this study are derivatives of the serotype O:8 strain WA314 harbouring the virulence plasmid pYVO8 (Heesemann & Laufs, 1983). WAC is the plasmidless derivative of WA314. WAC(pT3SS) is a derivative of WAC, which harbours the pT3SS plasmid encoding for the type three secretion system (T3SS). WA314ΔYopM, WA314ΔYopQ, WA314ΔYopH, WA314ΔYopO, WA314ΔYopT and WA314ΔYopE were constructed by replacing the corresponding gene in WA314 with a kanamycin resistance cassette (Trülzsch et al., 2004). WA314ΔYopP was generated by insertional inactivation of yopP gene (Ruckdeschel et al., 2001). WA314ΔYopMP and WA314ΔYopPQ are derivatives of *Yersinia enterocolitica* WA314. WA314ΔYopMQ and WA314 ΔYopMPQ were generated using a CRISPR-Cas12a-assisted recombineering approach like described previously (Rudolph et al., 2022; Yan et al., 2017). Table S1 provides an overview of the bacterial strains used in this study.

### Cell culture

Human peripheral blood monocytes were isolated from buffy coats as described in Kopp et al., 2006 (Kopp et al., 2006). Cells were cultured in Monocyte medium (RPMI1640, 20 % autologous serum, 1 % Penicillin/Streptomycin) at 37 °C and 5 % CO_2_. The medium was changed every three days until cells were differentiated into macrophages on day 7 after isolation. Macrophages were used for infection 1 week after the isolation except for RNA-seq samples from batch 1 in Table S2 for which cells were used after 2 weeks of isolation. Gene expression profiles were not affected by the time after isolation (Bekere et al., 2021).

### Infection of cells

On the day before infection of primary human macrophages the cell medium was changed to RPMI1640 without antibiotics and serum and precultures of *Y. enterocolitica* strains (Table S1) were grown overnight in LB medium with appropriate antibiotics at 27 °C and 200 x rpm. On the day of infection, precultures were diluted 1:20 in fresh LB medium without antibiotics and incubated for 90 min at 37 °C and 200 x rpm to induce activation of the *Yersinia* T3SS machinery and Yop expression. Afterwards bacteria were pelleted by centrifugation for 10 min at 6.000 x g, 4 °C and resuspended in 1 ml ice-cold PBS containing 1 mM MgCl_2_ and CaCl_2_. The optical density OD_600_ was adjusted to 3.6 and afterwards macrophages were infected at multiplicity-of-infection (MOI) from 100 to 500 as indicated in the figure legends. Cell culture plates were centrifuged for 2 min at RT and 200 x g to sediment bacteria on the cells and synchronize infection. Cells were incubated at 37 °C for 1.5 h to 6 h. For experiments using inhibitors for NF-κB pathway (TPCA, Cell Signaling) and MAPK pathway (PD98059, Cell Signaling & SB203580, Cayman Chemical), inhibitors were added 30 - 60 min prior infection at a final concentration of 10 µM.

### Histone extraction

Histones were extracted from primary human macrophages following modified acid extraction protocol from Shechter et al. (Shechter et al., 2007). 0,5 - 1,5 * 10^6^ primary human macrophages were washed with ice-cold PBS and harvested in 1 ml PBS. Cells were pelleted by centrifugation at 700 x g for 5 min at 4 °C and the supernatant was discarded. Lysis of the cells was done by incubation of the pellet in 0.5 ml Hypotonic lysis buffer (10 mM Tris-HCl, pH 8.0, 1mM KCl, 1mM DTT, 1.5 mM MgCl_2_, 1 x Complete Protease inhibitor, 1 x PhosStop Phosphatase Inhibitor) rotating at 4 °C for 60 - 90 min and controlled by microscopy. Isolated nuclei were pelleted by centrifugation at 10.000 x g for 10 min at 4 °C, supernatant was discarded and the pellet was resuspended in 0.2 M HCl and lysed by incubating at 4 °C rotating overnight. Nuclear debris was removed by centrifugation at 16.000 x g for 10 min at 4 °C and the supernatant was transferred to a new tube. Histones were precipitated using incubation with trichloric acid (TCA) (final concentration 33 %) for 30 min on ice. Pelleting of precipitated histones was carried out by centrifugation at 16.000 x g for 10 min at 4 °C and the supernatant was discarded. The pellet was washed twice with 500 µl ice cold acetone, centrifuged in between at 16.000 x g for 5 min at 4 °C, finally air dried and dissolved in 100 µl millipore water. Concentration of the dissolved histones was determined using Bradford protein assay and the quality and purity of the extracted histones was controlled by SDS-PAGE and subsequent Coomassie staining of the gel.

### Western blot analysis

Equal amounts of histone proteins were separated by SDS-PAGE and transferred to polyvinylidene difluoride (PVDF) membrane (Immobilon-P, Millipore, Schwalbach, Germany) by semi-dry blotting. The membrane was incubated with 3 % Bovine serum albumin (BSA) (w/v) in TBS supplemented with 0.05 % Tween 20 (TBS-T) for 30 - 60 min and subsequently with primary antibodies at 4 °C overnight. Primary antibodies used in this study were rabbit anti-Histone H3 antibody (Cell signaling, 1:3000 diluted), rabbit anti-Histone H3S10ph (Invitrogen and Abcam, 1:1000 diluted), rabbit anti-Histone H3K9me3 (Abcam, 1:1000 diluted) and rabbit anti-Histone H3K9me3S10ph (Abcam, 1:1000 diluted). The membrane was washed thrice with TBS-T and afterwards incubated at room temperature for 1 h with donkey anti-rabbit IgG (Cell Signaling, GE Healthcare) as a secondary antibody in a dilution of 1:10.000 - 1:25.000. Membrane was washed thrice with TBS-T and antibody signals were visualized with chemiluminescence technology (Supersignal West Femto, Pierce Chemical, Rockford, USA) and captured on X-ray films (Fujifilm, Düsseldorf, Germany). Developed films were scanned (CanonScan 4400 F, Canon, Tokio, Japan) to quantify protein band intensity. Quantification of signal intensity of scanned films was performed using ImageJ analysis software Version 1.53 (National Institute of Health, NIH).

### Immunofluorescence staining and confocal microscopy

Primary human macrophages were washed once with PBS and detached with Accutase (Thermo Fisher Scientific, Waltham, USA) for 30 min at 37 °C. Cells were scrapped off, mixed with equal volume of RPMI1640 medium (Gibco, Carlsbad, USA), counted and seeded in a density of 6 * 10^4^ cells onto glass coverslips (Marienfeld GmbH, Lauda-Königshafen, Germany). After 30 min at 37 °C, the medium was changed to Monocyte medium (RPMI1640, 20 % autologous human serum, 1 % Penicillin/Streptomycin) and cells were incubated overnight at 37 °C. Infection was done as described above. After infection, cells on coverslips were washed once with PBS, fixed with 4 % PFA in PBS for 7 min and washed twice with PBS. Cells were permeabilized with 0.1 % Triton X-100 in PBS for 15 min at RT, washed twice afterwards and blocked with blocking solution (PBS, 3 % (w/v) BSA, 0.05 % Triton X-100) for 30 - 60 min at RT in a humid chamber. Next coverslips were transferred into blocking solution containing the desired primary antibodies (mouse anti-ASC, Santa Cruz, 1:50; rabbit anti-Caspase-1, Invitrogen, 1:100) and incubated overnight at 4 °C. Washing with PBS was done thrice and incubation with secondary antibodies, DAPI and Phalloidin coupled to Alexa Fluor dyes diluted 1:200 in blocking solution was done for 60 min at RT in a humid chamber. Coverslips were washed thrice finally and mounted in Prolong Glass Antifade Mountant on object slide. Images were acquired using the laser scanning microscope Olympus FV 3000 with a 20x air objective (NA 0.8) or the 60x oil objective (NA 1.4) and the Olympus FV3000 Software (Evident Europe GmbH, Germany).

### Ca^2+^-imaging of primary human macrophages during *Yersinia enterocolitica* infection

Primary human macrophages were differentiated as described above. For Ca^2+^ imaging cells were seeded in 8-well chamber slides (ibidi) and loaded with Cal520-AM (5 µM), FuraRed-AM (10 µM) and Pluronic F127 (0.05 %) in RPMI for 60 min at 37 °C. Cells were washed and maintained in Ca^2+^-measuring buffer (NaCl 140 mM, KCl 5 mM, MgSO_4_ 1 mM, CaCl_2_ 1 mM, NaH_2_PO4 1 mM, Glucose 5.5 mM, HEPES 20 mM, pH 7.4) (Wolf et al., 2015). Imaging was performed with the spinning disk microscope Visitron SD-TIRF (Nikon Eclipse TiE, Nikon) with a 20 x CFI Plan Fluor DLL Phase objective (NA 0.51) and a sCMOS camera (Photometrics Prime 95B) in an incubation unit at 37 °C and 5.0 % CO_2_. Acquisition of images was carried out using the VisiView v4 software (Visitron Systems). The following settings were used for imaging Ca^2+^-signals: excitation (ex): 488 nm; emission1 (em1): 525/50 nm; em2: 700/75 nm; exposure time: 150 ms; acquisition rate: 3 frames per minute. Bacteria were prepared as described above and kept in Ca^2+^-measuring buffer. Cells were infected with a MOI of 50 3 min after the start of acquisition to capture baseline fluorescence. Imaging was conducted continuously for a period of 120 min after infection (Smail et al., 2018). Cells were stimulated with 1 µM Thapsigargin as a positive control 120 min following infection. Calibration of R_max_ (10 µM Ionomycin) and R_min_ (10 µM Ionomycin and 10 mM EGTA) was performed after each experiment (Wolf et al., 2015). Images were background and motion corrected (Dubbs et al., 2016). Single cells were defined as regions of interest (ROI). Image processing was performed using Fiji (Schindelin et al., 2012). Intracellular Ca^2+^-concentration was calculated with the *K*d of Cal520-AM (*K*d = 320 nM) (Grynkiewicz et al., 1985). Ca^2+^ spikes were defined as increase of ΔF_em1_> 20%, which exceeded the standard deviation of mock-infected macrophages by more than threefold (Chang-Graham et al., 2020). Quantification of fluorescence intensity was performed using Microsoft Excel, MATLAB R2023b (MathWorks) and with the assistance of ChatGPT (OpenAI).

### Statistical analysis

Statistical analysis was performed using Graph pad prism version 10.0. At least three independent experiments were compared by paired t-test, one-way ANOVA with Bonferronís post-test or two-way ANOVA if not indicated otherwise. p-values ≤ 0.05 were considered statistically significant.

### RNA-seq

Total RNA of 1-2 x 10^6^ human macrophages (for number of biological replicates/ macrophage donors per condition, see Table S2) was isolated using RNeasy extraction kit (Qiagen) including DNase treatment according to manufacturer’s instructions. RNA integrity of the isolated RNA was analyzed with the RNA 6000 Nano Chip (Agilent Technologies) on an Agilent 2100 Bioanalyzer (Agilent Technologies). mRNA was extracted using the NEBNext Poly(A) mRNA Magnetic Isolation module (New England Biolabs) and RNA-seq libraries were generated using the NEBNext Ultra RNA Library Prep Kit for Illumina (New England Biolabs) as per manufacturer’s recommendations. Concentrations of all samples were measured with a Qubit 2.0 Fluorometer (Thermo Fisher Scientific) and fragment lengths distribution of the final libraries was analyzed with the DNA High Sensitivity Chip (Agilent Technologies) on an Agilent 2100 Bioanalyzer (Agilent Technologies). All samples were normalized to 2 nM and pooled equimolar. The library pool was sequenced on the NextSeq500 (Illumina) with 1 x 75 bp for batch 2 samples, 1 x 50 bp for batch 1 samples and 1 x 75 bp for batch 3 samples (Table S2).

### RNA-seq data

Part of the RNA-seq data were obtained from already publicly available sources: ArrayExpress database at EMBL-EBI (www.ebi.ac.uk/arrayexpress) under accession number E-MTAB-10473 and European Nucleotide Archive (ENA) at http://www.ebi.ac.uk/ena/data/view/PRJEB10086. RNA-seq data for the first time used in this study have been deposited in the ArrayExpress database at EMBL-EBI (www.ebi.ac.uk/arrayexpress) under accession number E-MTAB-10602 and E-MTAB-15035. For detailed description of sample sources and batches used in this study refer to Table S2.

### RNA-seq analysis

Raw FASTQ files from RNA sequencing were processed using the nf-core/rnaseq pipeline (v1.3) (https://nf-co.re/rnaseq/) within the nf-core framework (Ewels et al., 2020). The analysis was performed with Nextflow (v21.04.0) (Di Tommaso et al., 2017), utilizing STAR (v2.6.1d) (Dobin et al., 2013) for alignment to the GRCh38 human genome. Gene-level quantification was carried out using featureCounts (v2.0.1) (Liao et al., 2014), which counts reads mapped to each gene. Transcript levels counts were imported using tximport (v1.30.0) (Soneson et al., 2015) package and summarized to gene levels counts with EnsDb.Hsapiens.v86 package (DOI: 10.18129/B9.bioc.EnsDb.Hsapiens.v86) and used for differential expression analysis with DESeq2 (v1.42.1) (Love et al., 2014) package. For differential expression analysis design formula “design = ∼batch+Condition” to correct for batch effect. For plotting of PCA plots and heatmaps counts were transformed using variance stabilized transformation followed by removal of batch effect using limma (v3.58.1) package (Law et al., 2014; Ritchie et al., 2015). Differentially expressed genes were defined as log2 fold change ≤ −2 or ≥ +2 and adjusted p value ≤ 0.01. For analyzing DEGs for naïve vs LPS-stimulated macrophages from publicly available datasets (Novakovic et al., 2016; Park et al., 2017) DEGs were defined as log2 fold change ≤ −1 or ≥ +1 and adjusted p value ≤ 0.05. DEG analysis of RNA-seq are found in Table S3. Pathway analysis was performed using DAVID analysis tool (Huang da et al., 2009).

For analysis of counterregulated genes between WA314ΔYopQ, WA314ΔYopM and WA314ΔYopP DEGs between WAC and WA314 6h were used and further selected if the same genes were upregulated by WA314ΔYopP vs WA314 6h (log2 fold change ≥ +1) but downregulated by WA314ΔYopM or WA314ΔYopQ vs WA314 6h (log2 fold change ≤ −1) or if the same genes were downregulated by WA314ΔYopP vs WA314 6h (log2 fold change ≤ −1) but upregulated by WA314ΔYopM or WA314ΔYopQ vs WA314 6h (log2 fold change ≥ +1).

### TF motif analysis

TF motif enrichment for known motifs was performed using HOMER package (Heinz et al. 2010). Command *finδMotifs.pl* was used and a list of gene symbols was supplied as an input. Motifs were searched in the region 400 bp upstream and 100 bp downstream of the TSS by specifying parameters *-start −400 -end 100*. For presentation of enriched TF motifs results from known motifs were used.

### Boxplots

Boxplots were generated using ggplot2 (3.5.1) in RStudio (2024.09.1+394). Boxes encompass the twenty-fifth to seventy-fifth percentile changes. Whiskers extend to the tenth and ninetieth percentiles. Outliers are depicted with black dots. The central horizontal bar indicates the median.

### Chromatin immunoprecipitation and sequencing (ChIP-seq)

ChIP-seq with formaldehyde crosslinking was performed as described previously (Bekere et al., 2021). Macrophages (3–10 x 10^6^ cells per condition, 2–3 biological replicates/ macrophage donors per condition) were washed once with warm PBS and incubated for 30 min at 37°C with accutase (eBioscience) to detach the cells. For ChIP BSA-blocked ChIP grade protein A/G magnetic beads (Thermo Fisher Scientific) were added to the chromatin and antibody mixture and incubated for 2 h at 4°C rotating to bind chromatin-antibody complexes. Samples were incubated for ∼3 min with a magnetic stand to ensure attachment of beads to the magnet and mixed by pipetting during the wash steps. Eluted DNA was subjected to ChIP-seq library preparation. Input chromatin DNA was prepared from 1/4 of chromatin amount used for ChIP. Antibodies used for ChIP were anti-H3S10ph (Invitrogen, 9H12L10, 6 μl (3 μg) per ChIP), anti-H3K9me3S10ph (abcam, ab5819, 2 μg per ChIP).

ChIP-seq libraries were constructed with 1–10 ng of ChIP DNA or input control as a starting material. Libraries were generated using the NEXTflex ChIP-Seq Kit (Bioo Scientific) as per manufacturer’s recommendations. Concentrations of all samples were measured with a Qubit Fluorometer (Thermo Fisher Scientific) and fragment length distribution of the final libraries was analyzed with the DNA High Sensitivity Chip on an Agilent 2100 Bioanalyzer (Agilent Technologies). All samples were normalized to 2 nM and pooled equimolar. The library pool was sequenced on the NextSeq500 (Illumina) with 1 x 75 bp and total approx. 20 to 40 million reads per sample.

### ChIP-seq analysis

Raw FASTQ files from ChIP sequencing were processed using the nf-core/chipseq pipeline (v1.0.0) (https://nf-co.re/chipseq/) within the nf-core framework (Ewels et al., 2020). The pipeline was executed with Nextflow v0.26.4 (Di Tommaso et al., 2017), starting with TrimGalore (http://www.bioinformatics.babraham.ac.uk/projects/trim_galore/) to trim low-quality bases (Phred score cutoff of 20) and adapter sequences. The trimmed reads were then aligned to the GRCh37 human genome using BWA (v0.7.12) (Li & Durbin, 2009), generating an alignment format. Samtools (Li et al., 2009) was used to perform manipulations on the alignment format, such as sorting, indexing, and format conversions. To remove duplicate reads, Picard (https://broadinstitute.github.io/picard/) was applied, and BEDTools (Quinlan & Hall, 2010) was used to generate BED files from the aligned data.

For defining H3K9me3 regions in macrophages from publicly available data (Novakovic et al., 2016), SICER2 (Zang et al., 2009) was used using input ChIP-seq data as a control. Peaks were filtered to exclude blacklisted regions (Amemiya et al., 2019) and regions with p value larger than 0.05 and regions with fold change less than 2. ChIP-seq signal was plotted using EaSeq (Lerdrup et al., 2016) (https://easeq.net) and IGV (Robinson et al., 2011).

For analysis of chromatin states using ChromHMM analysis (Ernst & Kellis, 2012, 2017) bed files were binarized using bin size of 1000bp. Different number of emissions (chromatin states) was tested and seven emissions were selected to depict all unique chromatin states.

ChIP-seq data have been deposited in the ArrayExpress database at EMBL-EBI (www.ebi.ac.uk/arrayexpress) under accession number E-MTAB-14995.

## Acknowledgements

We thank Daniela Indenbirken for generating the RNA-seq libraries and Frank Bentzien (UKE Transfusion Medicine) for buffy coats. This work was supported by the research consortium EPILOG, funded by the Stiftung zur Förderung der wissenschaftlichen Forschung in Hamburg. S. Kulnik and M. Aepfelbacher were funded by DFG RTG2771. We thank UKE Microscopy Imaging Facility (UMIF) for training and support. The laser scanning microscope Olympus FV 3000 was funded by the DFG (code: INST 152/933-1)

---

**Figure S1:**
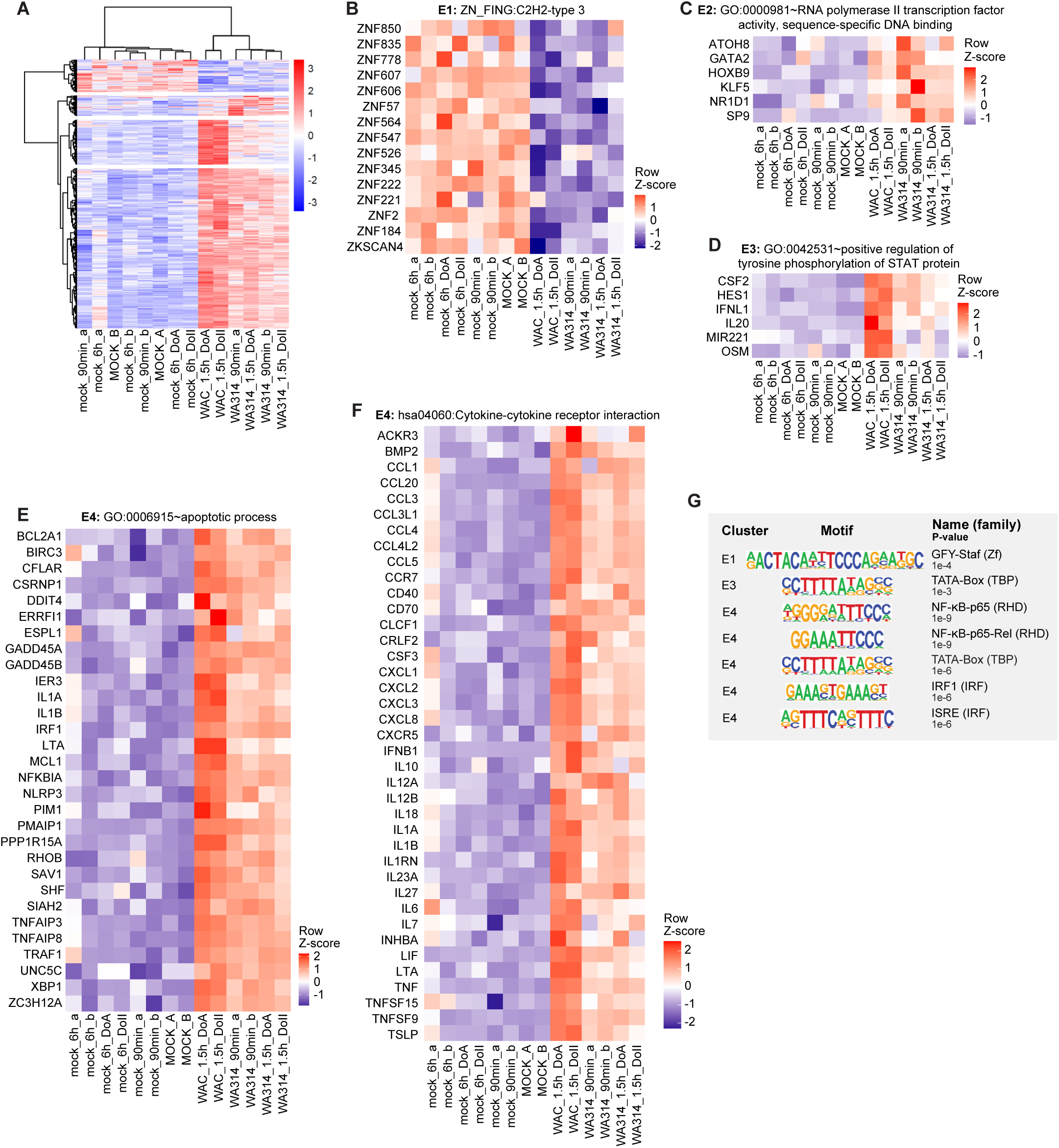
A: Clustered Heatmap of all DEGs for comparisons between mock, WAC and WA314 (also named WAP) infected macrophages after 1.5 h of infection. Clustering showed 4 major clusters. Gene vsd counts were row-scaled (row Z-score). All replicates used in this study are shown Table S2). B: Heatmap of row scaled RNA-seq vsd counts for genes belonging to Zinc finger genes from the C2H2 type 3 enriched in cluster E1. C: Heatmap of row scaled RNA-seq vsd counts for genes belonging to the RNA polymerase II transcription factor activity, sequence-specific DNA binding pathway enriched in cluster E2. D: Heatmap of row scaled RNA-seq vsd counts for genes belonging to the positive regulation of tyrosine phosphorylation of STAT protein pathway enriched in cluster E3. E: Heatmap of row scaled RNA-seq vsd counts for genes belonging to the apoptotic process pathway enriched in cluster E4. F: Heatmap of row scaled RNA-seq vsd counts for genes belonging to the cytokine-cytokine receptor interaction pathway enriched in cluster E4. G: Representative enriched transcription factor motifs in genes from clusters E1 - E4 (Fig 2A-C). Vsd counts for generation of heatmaps can be found in the Table S4.

**Figure S2:**
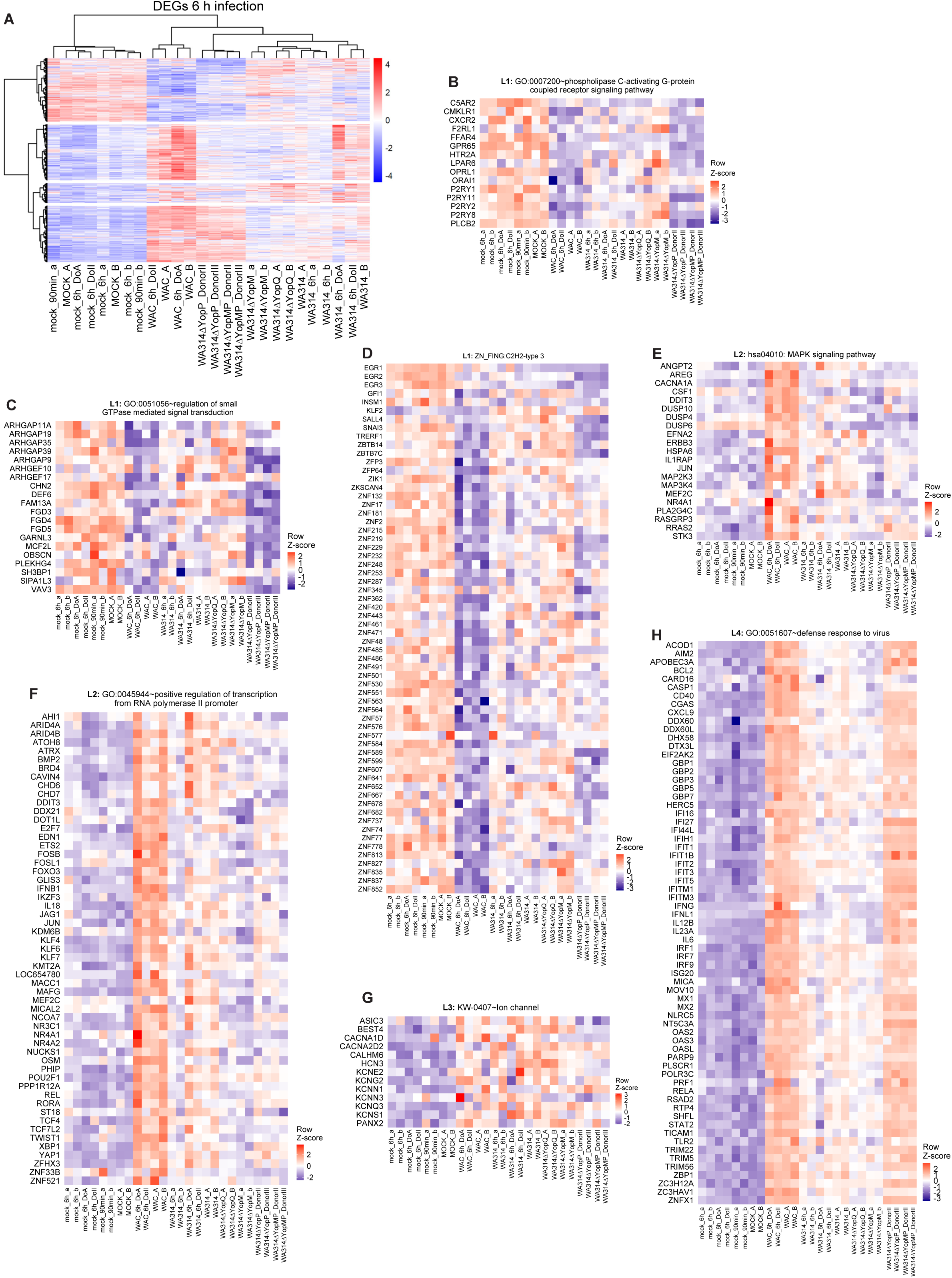
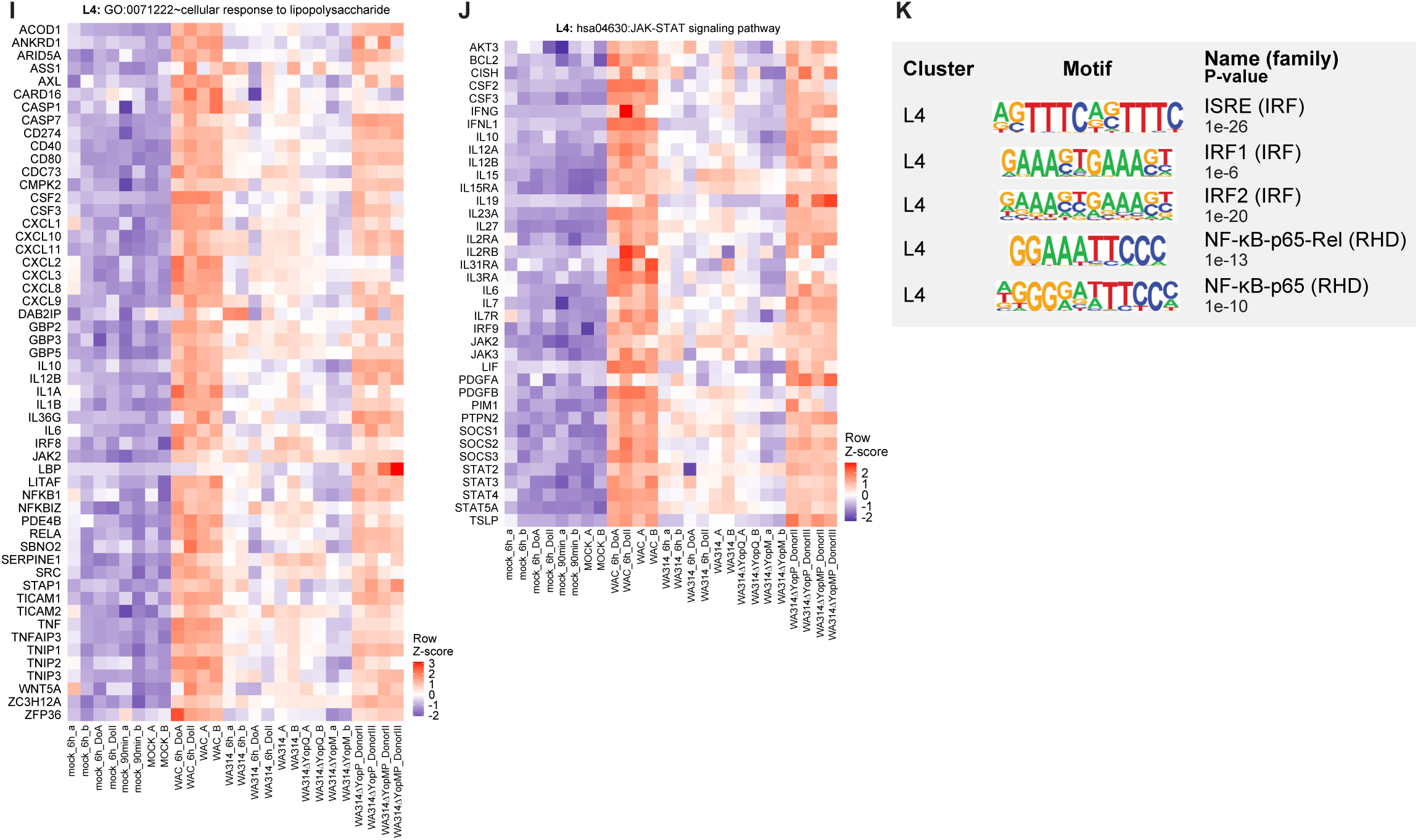
A: Clustered Heatmap of all DEGs for comparisons between mock, WAC, WA314, WA314ΔYopP, WA314ΔYopMP, WA314ΔYopM, WA314ΔYopQ infected macrophages after 6 h of infection. Clustering showed 4 major clusters. Gene vsd counts were row-scaled (row Z-score). All replicates used in this study are shown. B: Heatmap of row scaled RNA-seq vsd counts for genes belonging to the phospholipase C-activating G-protein coupled receptor signaling pathway enriched in cluster L1. C: Heatmap of row scaled RNA-seq vsd counts for genes belonging to the regulation of small GTPase mediated signal transduction pathway enriched in cluster L1 from all replicates used in this study. D: Heatmap of row scaled RNA-seq vsd counts for genes belonging to Zinc finger genes from the C2H2 type 3 enriched in cluster L1 from all replicates used in this study. E: Heatmap of row scaled RNA-seq vsd counts for genes belonging to MAPK signaling pathway enriched in cluster L2 from all replicates used in this study. F: Heatmap of row scaled RNA-seq vsd counts for genes belonging to the positive regulation of transcription from RNA polymerase II promotor pathway enriched in cluster L2 from all replicates used in this study. G: Heatmap of row scaled RNA-seq vsd counts for genes belonging to the group of ion channels enriched in cluster L3 from all replicates used in this study. H: Heatmap of row scaled RNA-seq vsd counts for genes belonging to defence response to virus pathway enriched in cluster L4 from all replicates used in this study. I: Heatmap of row scaled RNA-seq vsd counts for genes belonging to the cellular response to lipopolysaccharide pathway enriched in cluster L4 from all replicates used in this study. J: Heatmap of row scaled RNA-seq vsd counts for genes belonging to the JAK-STAT signaling pathway enriched in cluster L4 from all replicates used in this study. K: Representative enriched transcription factor motifs in genes from L1 - L4 clusters in Fig 3E. Vsd counts for generation of heatmaps can be found in the Table S4.

**Figure S3:**
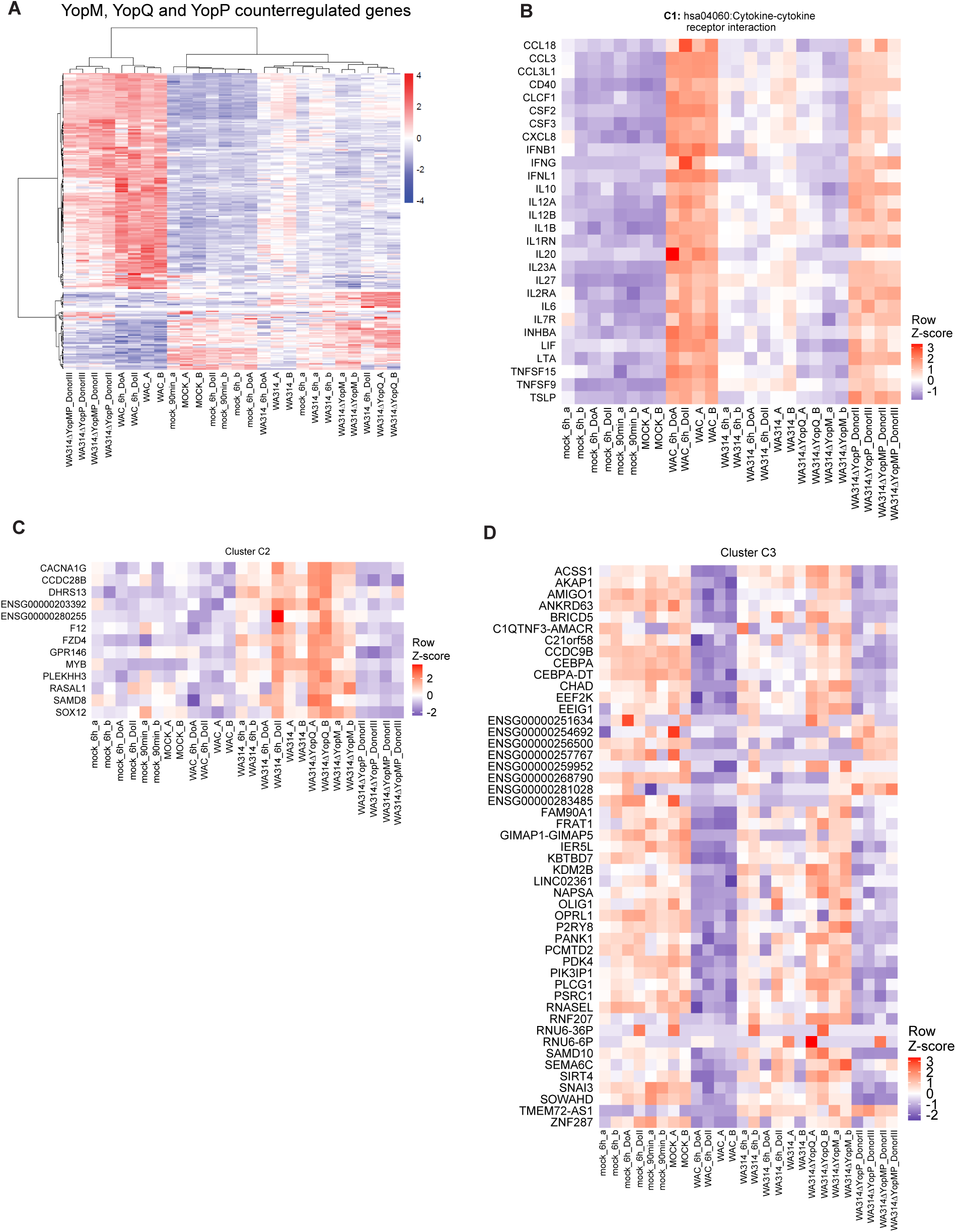
A: Clustered heatmap of genes counterregulated by YopM, YopP and YopQ in comparisons between WAC- and WA314-infected macrophages (6 h). Gene vsd counts were row-scaled (row Z-score) and clustering identified 3 major clusters. All replicates used in this study are shown. B: Heatmap of row scaled RNA-seq vsd counts for genes belonging to cytokine-cytokine receptor interaction pathway enriched in cluster C1 from all replicates used in this study. C: Heatmap of row scaled vsd counts for genes in cluster C2 showing all replicates used in the study. D: Heatmap of row scaled vsd counts for genes in cluster C3 showing all replicates used in the study. Vsd counts for generation of heatmaps can be found in the Table S4.

**Figure S4:**
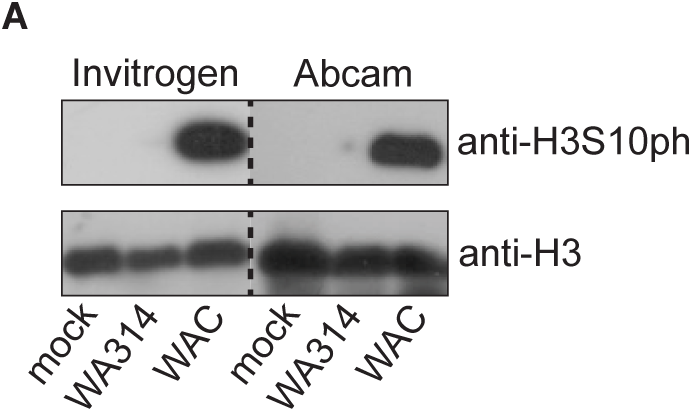
A: Western blot showing H3S10ph with two different anti-H3S10ph antibodies and H3 in macrophages not infected (mock) or infected with WA314 or WAC for 3 h with MOI of 100. H3 bands serve as loading control.

**Figure S5:**
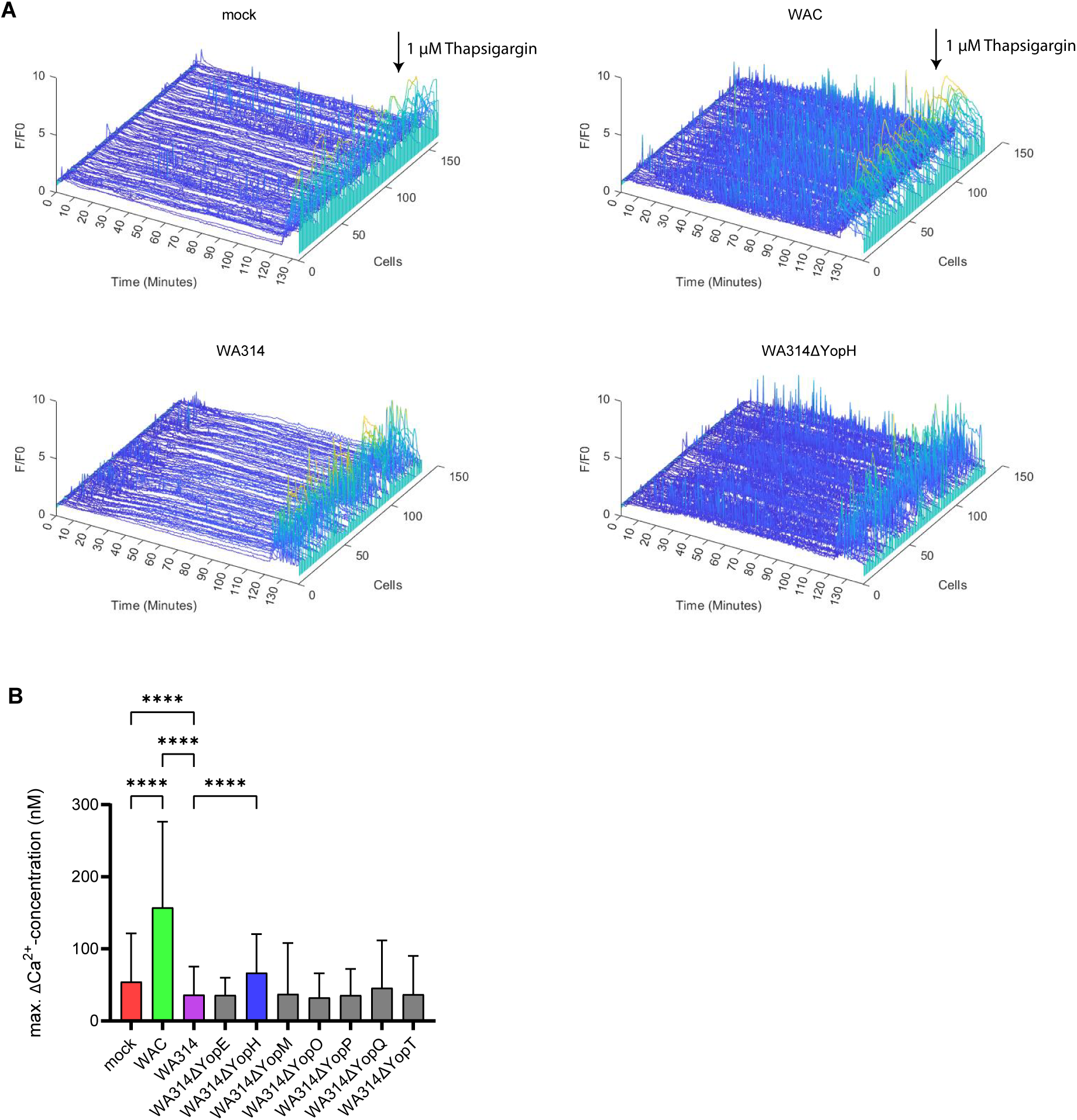
A: Representative normalized (F_em1_/F_em1_0) single-cell fluorescence intensity profiles. Human macrophages were not infected (mock) or infected with indicated strains with a MOI of 50 for 2h and loaded with Cal520-AM and FuraRed-AM. Cells were imaged with 3 frames per min (fpm) for 120 min. Stimulation with 1 µM Thapsigargin at 120 min post infection (mpi) was used as a positive control. B: Quantification of maximal ΔCa^2+^-concentration/cell.

**Movie S1:**

Representative videos (F_em1,_ 3 fpm) of human macrophages not infected (mock) or infected with indicated strains with a MOI of 50 for 2h and loaded with Cal520-AM and FuraRed-AM to visualize intracellular Ca^2+^ fluctuations. Stimulation with 1 µM Thapsigargin at 120 mpi was used as a positive control. Scale bar: 100 µm. Calibration bar: fluorescence intensity (F_em1_) (a.u.). Time stamper: mm:ss.

